# Practical considerations for using K3 cameras in CDS mode for high-resolution and high-throughput single particle cryo-EM

**DOI:** 10.1101/2020.11.08.372763

**Authors:** Ming Sun, Caleigh Azumaya, Eric Tse, Adam Frost, Daniel Southworth, Kliment A. Verba, Yifan Cheng, David A. Agard

## Abstract

Detector technology plays a pivotal role in high-resolution and high-throughput cryo-EM structure determination. Compared with the first-generation, single-electron counting direct detection camera (Gatan K2), the latest K3 camera is faster, larger, and now offers a correlated-double sampling mode (CDS). Importantly this results in a higher DQE and improved throughput compared to its predecessor. In this study, we focused on optimizing camera data collection parameters for daily use within a cryo-EM facility and explored the balance between throughput and resolution. In total, eight data sets of murine heavy-chain apoferritin were collected at different dose rates and magnifications, using 9-hole image shift data collection strategies. The performance of the camera was characterized by the quality of the resultant 3D reconstructions. Our results demonstrated that the Gatan K3 operating in CDS mode outperformed nonCDS mode in terms of reconstruction resolution in all tested conditions with 8 electrons per pixel per second being the optimal dose rate. At low magnification (64kx) we were able to achieve reconstruction resolutions of 149% of the physical Nyquist limit (1.8 Å with a 1.346 Å physical pixel). Low magnification allows more particles to be collected per image, aiding analysis of heterogeneous samples requiring large data sets. At moderate magnification (105kx, 0.834Å physical pixel size) we achieved a resolution of 1.65 Å within 9 hours of data collection, a condition optimal for achieving high-resolution on well behaved samples. Our results also show that for an optimal sample like apoferritin, one can achieve better than 2.5 Å resolution with 5 minutes of data collection. Together, our studies validate the most efficient ways of imaging protein complexes using the K3 direct detector and will greatly benefit the cryo-EM community.

## Introduction

Cryo-electron microscopy (cryo-EM), particularly single particle cryo-EM, has experienced tremendous success in its ability to provide structural information on how dynamic molecules perform their biological functions. Through technological advances in both hardware and software, it is now possible to determine structures of a wide range of biological complexes at or near atomic resolution using cryo-EM. Recently, the achievable resolution of single particle cryo-EM has been extended to well beyond 2Å, where individual atoms can be accurately localized in the cryo-EM density map (Nakane *et al*., 2020; Yip *et al*., 2020; Zhang *et al*., 2020). However, achieving such high resolution comes at a cost of time and resources, which will influence how broadly the technology will be employed to address fundamental biological questions.

Detector technology plays a pivotal role in high-resolution and high-throughput structure determination and drove the ‘resolution revolution’ in cryo-EM. The current direct detectors combine direct detection and single electron counting, which allows dramatic improvements in the Detective Quantum Efficiency (DQE) well beyond that previously possible. DQE is one of the most important quantitative metrics characterizing the performance of EM cameras and varies as a function of spatial frequency (McMullan *et al*., 2009, 2016; Li, Mooney *et al*., 2013; Mooney, 2007). Higher DQE values indicate less noise added by the detector itself to the final images. Therefore, maximizing DQE across all frequencies is a major objective during the detector design. During the course of detecting single electron events, the incident electron can distribute charge in adjacent pixels, allowing the incident location to be estimated to higher precision (known as super-resolution mode on Gatan cameras) (Li, Zheng *et al*., 2013; Booth *et al*., 2012). The 2x increased sampling reduces aliasing artifacts and provides the ability for usable information to go beyond the physical Nyquist.

The Gatan K3 detector (Ametek Gatan) further improves DQE compared to the previous generation K2 Summit detector by having improved image processing, a 4x higher internal frame rate and a higher pixel count. While many factors affect DQE in any single electron counting camera, for a given pixel design and sensitive layer thickness, two important ones are analog noise and frame rate. The former determines the smallest signal from an incoming electron that can be reliably counted, while the latter dictates the degree of systematic undercounting of events due to two electrons arriving within the frame readout interval (known as coincidence loss) ((Li, Zheng *et al*., 2013); Figure S1). The effects of coincidence loss can be visualized in noise power spectrum (NPS) curves (Figure S2; Table S1 and S2). For any given dose rate, the higher the internal camera frame rate the less coincidence loss and the higher the effective DQE. A practical trade-off is that choosing to work at lower dose rates translates into longer exposures and reduced throughput. Compared with the K2, the K3 also successfully implements a correlated double sampling (CDS) imaging mode which suppresses analog noise (McMullan *et al*., 2016) allowing a more reliable detection of weak events. This is accomplished by effectively subtracting a dark frame from each frame recorded, taking two read cycles. Thus, there is a dose-rate dependent trade-off between better detection in CDS mode vs. reduced coincidence loss in nonCDS mode.

To optimize the usage of the K3 for daily operation, we explored the balance between throughput and resolution and characterized the performance of the camera by evaluating the quality of 3D reconstructions from data collected at different dose rates and magnifications. Magnification determines the image pixel size (both physical and super-resolution) and Nyquist resolution limits. Since the DQE of a camera is a function of Nyquist resolution, altering the magnification places different spatial frequencies in different DQE ranges of the camera. At a given spatial resolution, higher magnification directly translates into improved DQE but comes at the expense of a reduced effective imaging area, limiting the number of particles that can be recorded within each image.

Apoferritin, a standard sample for evaluating data quality in cryo-EM (∼500 kDa with octahedral symmetry) was collected at 300kV on our standard X-FEG Titan Krios equipped with a Gatan BioQuantum energy filter and K3 camera. To maximize throughput, we utilized image shift (Mastronarde, 2005; Suloway *et al*., 2005) rather than stage motion to image nearby holes. In our collection scheme, a central point is localized using stage shift and images are shot in a regular 3×3 pattern of holes around the central location using a combination of beam tilt and image shift. At 64kx we are able to reach 1.8 Å, ∼50% higher than the 2.69 Å physical Nyquist frequency from 373,000 selected particles within 9 hours of image-shift data collection. This reconstruction far surpasses the physical Nyquist resolution in a standard data collection period in our facility. At 105kx, we reached 1.65 Å, 101% of the 1.67 Å physical Nyquist frequency, from 571,000 selected particles within 8 hours of data collection. In as short as 5 minutes, from 7,000 particles, apoferritin can be reconstructed to under 2.5 Å resolution using a 1.34 Å physical pixel size, under ideal conditions. In our higher magnification maps, we were able to discern high-resolution features including densities for hydrogens in the peptide backbone and certain amino acid side chains. Our results demonstrate that the Gatan K3 operating in CDS mode outperforms nonCDS mode in terms of reconstruction resolution in all tested conditions with 8 electrons per pixel per second (eps) being the most efficient dose rate to reach high-resolution.

## Results

### Imaging benchmark cryo-EM samples under different conditions on the Gatan K3 camera

In this study, we imaged apoferritin to systematically characterize the performance of the K3 camera with and without using CDS mode, as well as to elucidate the relationship between two important practical aspects of single particle cryo-EM: attainable resolution vs. throughput. Summarized in Table 1, we collected a total of eight datasets of apoferritin at different dose rates and magnifications. As described in detail in the methods section, all data sets were collected using the identical, whenever possible, or comparable microscope settings. The three most important variables we explored were magnification, dose rate, and CDS mode. Different dose rates were achieved by adjusting spot size without changing additional microscope parameters. Exposure times were adjusted so that the total electron dose per each image stack was held constant. Additionally, physical pixel sizes are 0.834 Å/pix at 105,000x magnification and 1.346 Å/pix at 64,000x magnification, which correspond to Nyquist limits of 1.668 Å and 2.692 Å, respectively. Although our analyses here focus on apoferritin, we also collected 4 data sets from the much smaller aldolase to confirm the generality of our observations (Table 1).

**Table 1.**
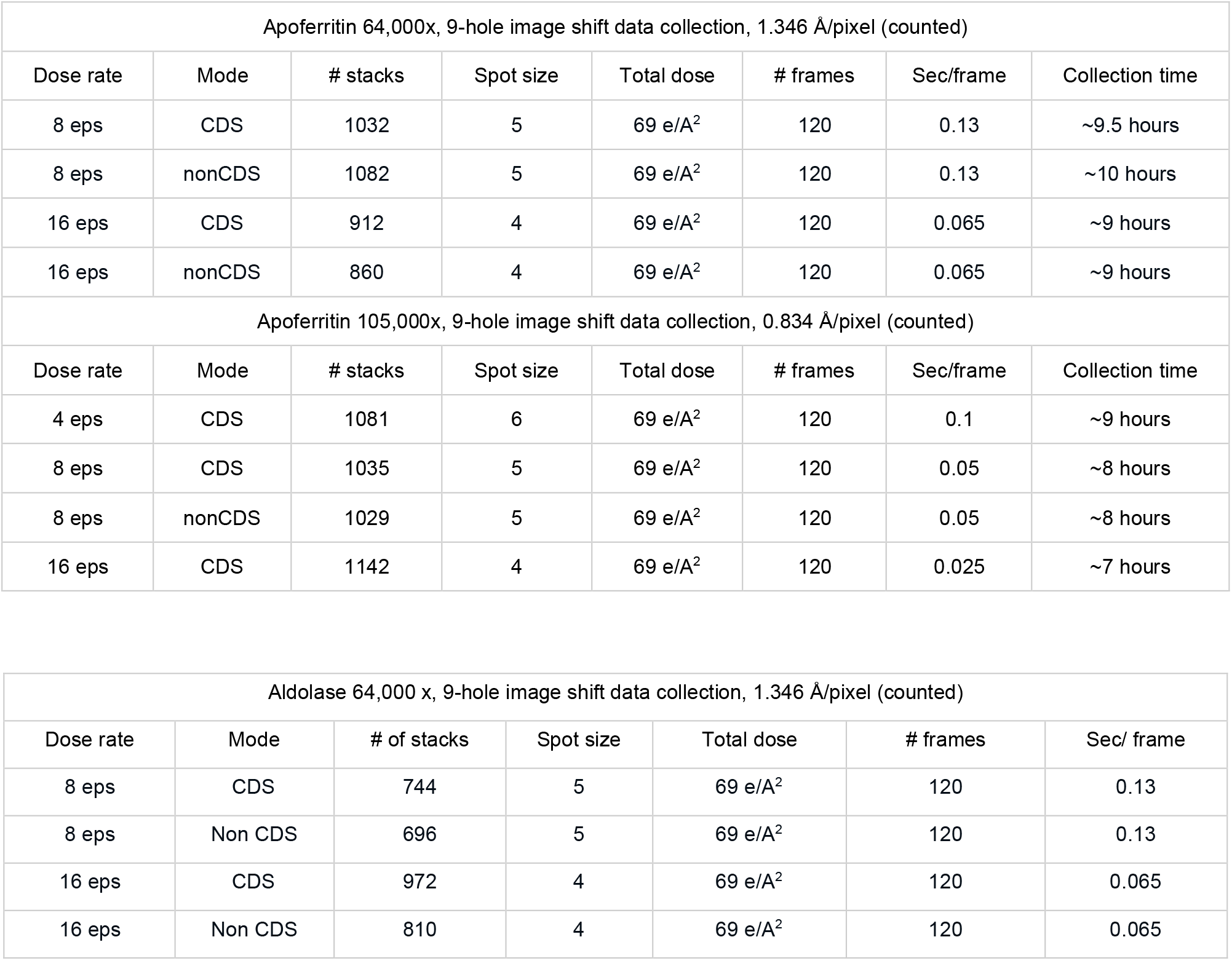
Data collection summary for apoferritin and aldolase data sets. All data sets were collected on a Titan Krios G3 at 300 kV equipped with a Gatan K3 camera and BioQuantum energy filter. Applicable units are denoted in the tables and data sets are separated by pixel size.

| Apoferritin 64,000x, 9-hole image shift data collection, 1.346 Å/pixel (counted) |  |  |  |  |  |  |  |
| --- | --- | --- | --- | --- | --- | --- | --- |
| Dose rate | Mode | # stacks | Spot size | Total dose | # frames | Sec/frame | Collection time |
| 8 eps | CDS | 1032 | 5 | 69 e/A<sup>2</sup> | 120 | 0.13 | ~9.5 hours |
| 8 eps | nonCDS | 1082 | 5 | 69 e/A<sup>2</sup> | 120 | 0.13 | ~10 hours |
| 16 eps | CDS | 912 | 4 | 69 e/A<sup>2</sup> | 120 | 0.065 | ~9 hours |
| 16 eps | nonCDS | 860 | 4 | 69 e/A<sup>2</sup> | 120 | 0.065 | ~9 hours |
| Apoferritin 105,000x, 9-hole image shift data collection, 0.834 Å/pixel (counted) |  |  |  |  |  |  |  |
| Dose rate | Mode | # stacks | Spot size | Total dose | # frames | Sec/frame | Collection time |
| 4 eps | CDS | 1081 | 6 | 69 e/A<sup>2</sup> | 120 | 0.1 | ~9 hours |
| 8 eps | CDS | 1035 | 5 | 69 e/A<sup>2</sup> | 120 | 0.05 | ~8 hours |
| 8 eps | nonCDS | 1029 | 5 | 69 e/A<sup>2</sup> | 120 | 0.05 | ~8 hours |
| 16 eps | CDS | 1142 | 4 | 69 e/A<sup>2</sup> | 120 | 0.025 | ~7 hours |

| Aldolase 64,000 x, 9-hole image shift data collection, 1.346 Å/pixel (counted) |  |  |  |  |  |  |
| --- | --- | --- | --- | --- | --- | --- |
| Dose rate | Mode | # of stacks | Spot size | Total dose | # frames | Sec/ frame |
| 8 eps | CDS | 744 | 5 | 69 e/A<sup>2</sup> | 120 | 0.13 |
| 8 eps | Non CDS | 696 | 5 | 69 e/A<sup>2</sup> | 120 | 0.13 |
| 16 eps | CDS | 972 | 4 | 69 e/A<sup>2</sup> | 120 | 0.065 |
| 16 eps | Non CDS | 810 | 4 | 69 e/A<sup>2</sup> | 120 | 0.065 |

### Influence of dose rates for imaging samples in CDS mode

We first explored the influence of dose rates on imaging frozen biological samples recorded in CDS mode through a series of experiments, ranging from 4eps to 16eps. The resolutions of 3D reconstructions determined from these data sets were evaluated in terms of both absolute value in angstroms and as a percentage of the corresponding physical Nyquist frequency limit (Figure 1A, Table S3). Comparing apoferritin data sets collected at 105kx, the one collected at a dose rate of 8eps reached a higher resolution than the ones at 4 and 16eps. The reconstruction from 200,000 particles yielded an average resolution of 1.75 Å, ∼95% of the physical Nyquist resolution, as compared to 1.78 Å from 4 and 16eps data sets. In addition, the benefit of collecting at 8 eps became even more pronounced when subsets of particles were used. A reconstruction from 10,000 particles reached 1.98 Å, ∼84% Nyquist frequency at 8eps, but only 2.16 Å, ∼77% (8% lower), at 16eps. Consistently, when comparing data subsets collected at a lower magnification, 64kx, 8eps data yielded higher resolutions than 16eps. However, these differences vanished when using 200,000 particles, for which both low magnification reconstructions reached 1.91 Å, an astonishing ∼141% of the physical Nyquist (Figure 1A, Table S3).

**Figure 1.**
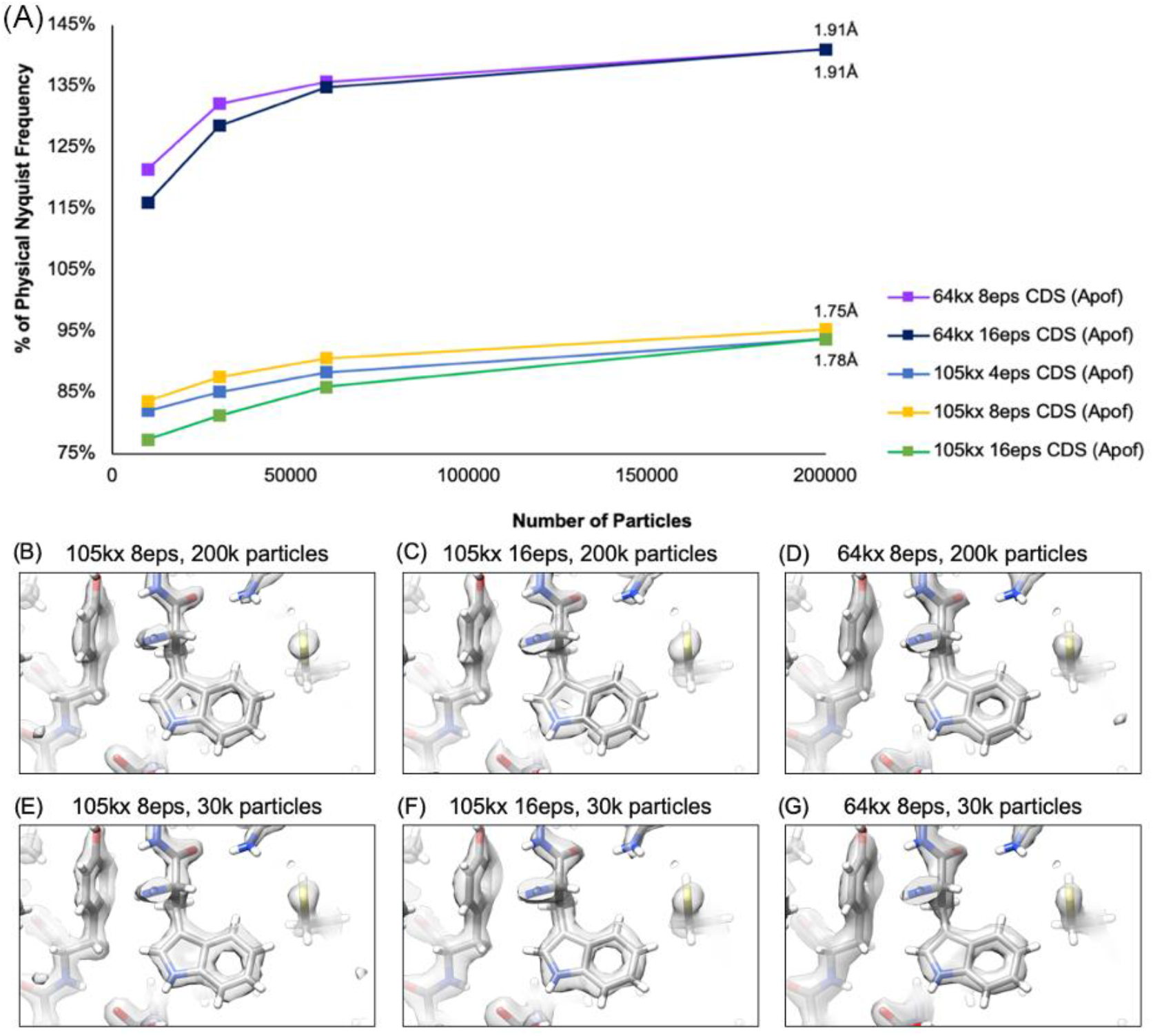
Influence of dose rate on reconstructions of apoferritin from data recorded on the K3 detector. **(A)** Results of 3D reconstructions from apoferritin data sets collected at different dose rates. The results are plotted as a function of percent of physical Nyquist frequency versus particle numbers in order to compare the effect of dose rate on data sets collected at different magnifications. 105kx correlates to a physical pixel size of 0.834 Å/pix and 64kx to 1.346Å/pix. The highest resolution achieved for each data set is labelled. **(B-D)** Cryo-EM maps calculated from 200k particles for **(B)** 8eps **(C)** 16eps data sets collected at 105kx magnification and **(D)** 8eps collected at 64kx magnification. **(E-G)** 3D reconstructions calculated from 30,000 particles for **(E)** 8eps and **(F)** 16eps data sets collected at 105kx magnification, and **(G)** 8eps collected at 64kx magnification. Data was processed in RELION v3.1, following standard protocols and sharpened using standard RELION post-processing protocols. These snapshots allow us to visualize the higher resolution structural information retained in the data collected at lower dose rates.

While the gains in resolution when using a dose rate of 8eps were modest in terms of the global FSC resolution estimate, there were clear improvements in the maps. We scrutinized the structural features of each amino acid, but focused here on the atomic details of Trp-93 (Figure 1B-G). At 8eps, the four distal carbons of the benzene ring can be resolved to an extent that the ring has a clear hexagonal shape, unlike in the 16eps map where it appears more circular. Also, the hole in the tryptophan pyrrole ring becomes clearer as dose rate is decreased. Consistent with the numerical resolution, the hole in the pyrrole ring is not resolved in the lower magnification reconstructions. Maps reconstructed from only 30,000 particles show similar quality trends to those observed with all the particles. For example, the density around the backbone carbonyl directly behind Trp-93 is notably better defined in the high magnification, 8eps map than in the other 30,000 particle maps.

### Influence of CDS mode under conditions of constant coincidence loss

Perhaps one of the most important new features on the K3 camera is the availability of CDS mode. To disentangle the conflicting influences of decreased noise and increased coincidence loss that are introduced when using CDS mode, we compared data collected such that the total coincidence losses, in principle, were held constant (Table S2). Specifically, apoferritin samples were exposed at 8eps CDS vs 16eps nonCDS at 64kx magnification, and 4eps CDS vs 8eps nonCDS at 105kx magnification.

**Figure 2** summarizes our results for apoferritin and shows that operating in CDS mode produced higher quality data and yielded higher resolution reconstructions at both magnifications. At 64kx magnification, the average resolution of apoferritin from 200k particles is 1.91 Å (141% of the Nyquist frequency) at 8eps in CDS mode, whereas at 16eps in nonCDS mode, the average resolution decreased by 7% to 1.98 Å. Smaller subsets of particles also reflect this trend, with the 8eps reconstructions improving ∼7-12% in terms of percentage of Nyquist frequency for each subset. At 105kx magnification, we observed ∼5% gain in resolution with 200k particles (Figure 2, Table S4). There is less separation between 4eps in CDS mode and 8eps in nonCDS mode at higher magnification, 3-5% for all subsets.

**Figure 2.**
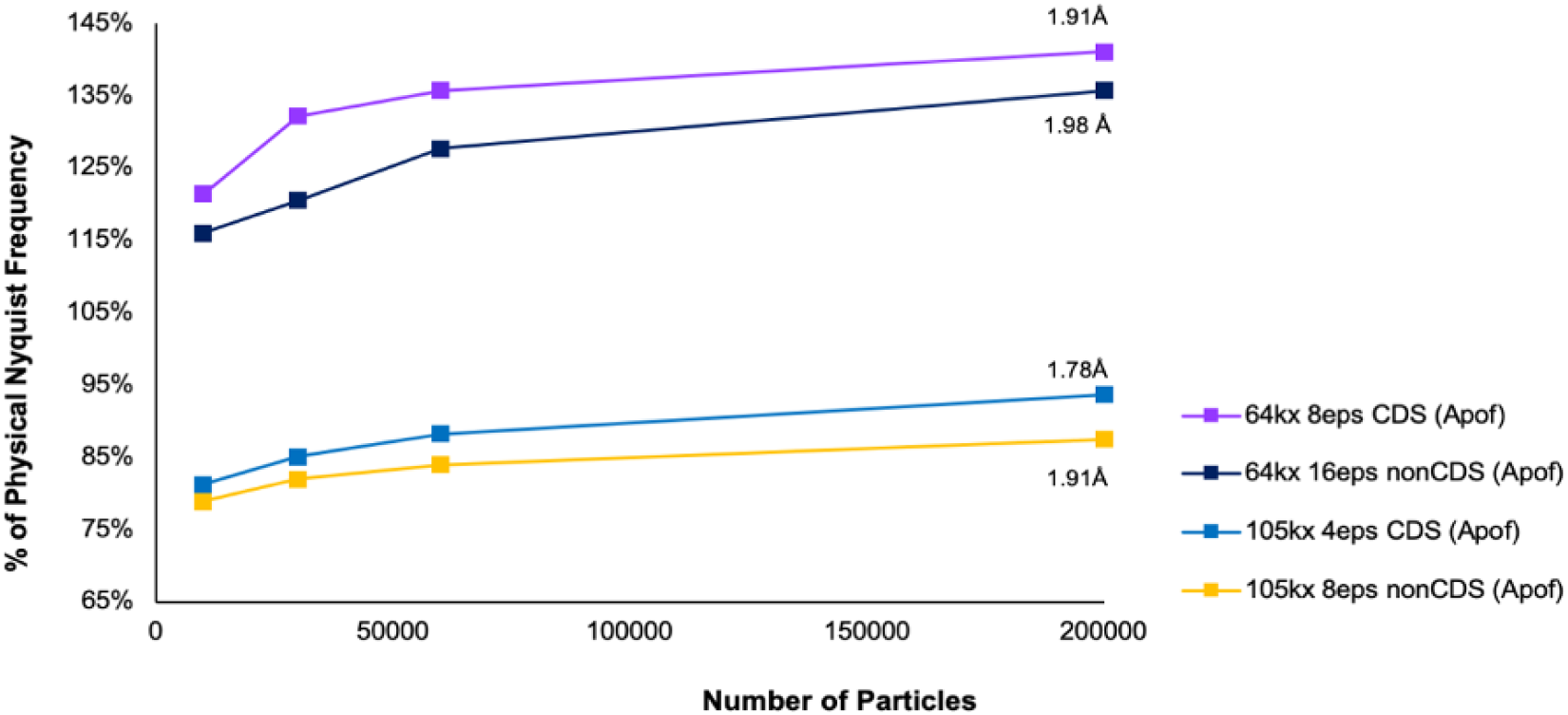
Benefits of imaging apoferritin in CDS mode when comparing data sets with equivalent coincidence loss of electron events. Comparing results of apoferritin in CDS and nonCDS modes where coincidence loss events are consistent. These results are plotted as a function of percent of physical Nyquist frequency versus particle number and highest achievable resolutions are labelled. Data was processed in RELION v3.1, following standard protocols (Scheres, 2012; Zivanov *et al*., 2018)

### Influence of CDS mode on imaging samples at constant dose rate

Under conditions of constant coincidence loss, the gain in using CDS was clear, however, this led to practical differences in exposure time. Next, we compared data collected at a constant dose rate, 8eps, in CDS and nonCDS mode. Again, comparison of the 3D reconstructions showed that gains from CDS mode compensated for increased coincidence losses at both magnifications, and that the benefit was most prominent at higher magnification (Figure 3, Table S5). At 105kx magnification, the achievable resolution with CDS at 8eps was 10% higher than without CDS, 1.75 Å compared to 1.94 Å when comparing reconstructions from 200,000 particles each. This trend held for all particle subsets sampled at high magnification, with differences from 5-10% of percentage Nyquist. At 64kx magnification, the gain in resolution when using CDS mode was only 2%. Compared with the gain from using CDS mode at constant coincidence loss (Figure 2), the gain at 105kx magnification from using CDS mode at constant dose rate was similar (Figure 3), while it was reduced for the 64kx magnification data sets.

**Figure 3.**
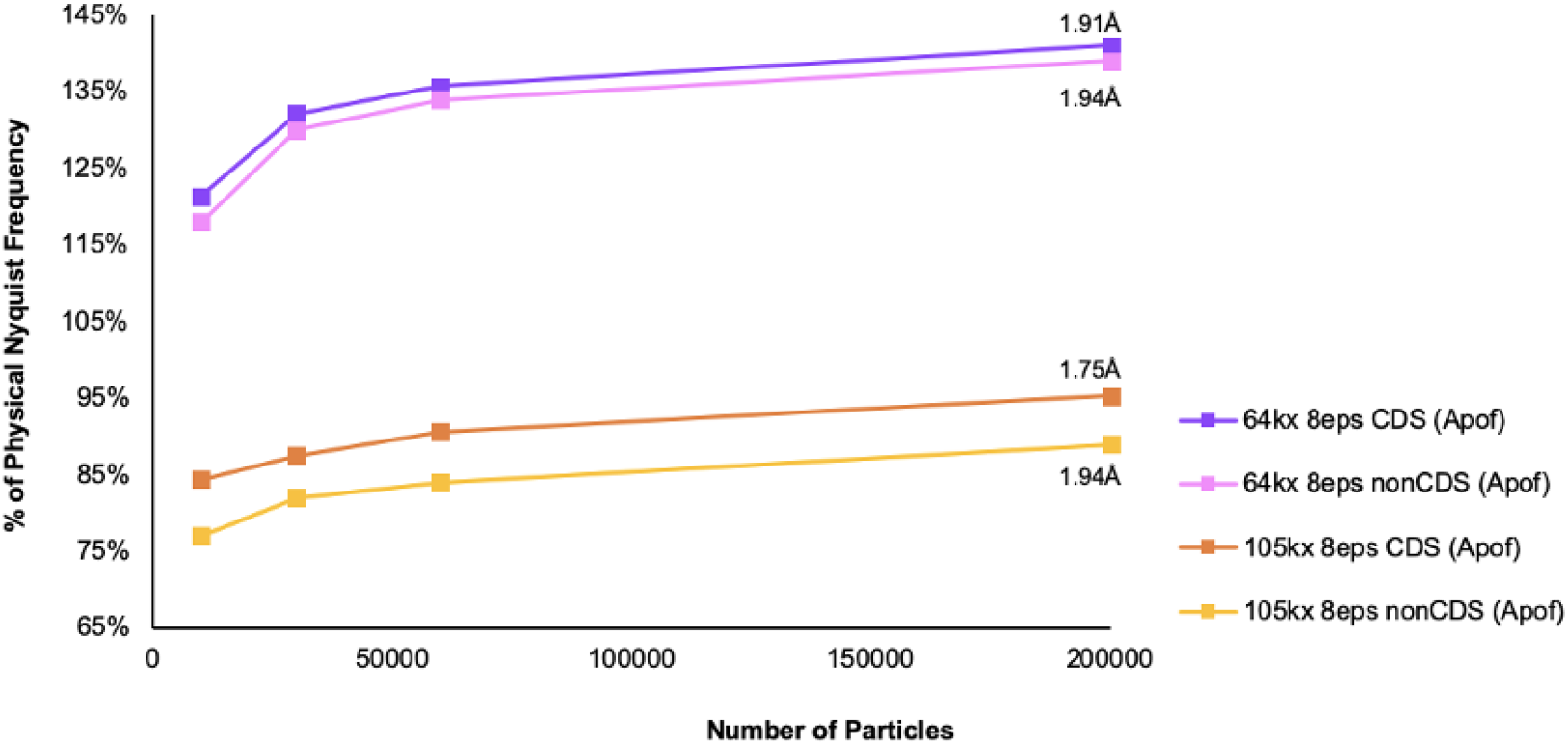
Practical consideration of imaging apoferritin using CDS mode at our recommended dose rate of 8eps. Comparing results of aldolase in CDS and nonCDS modes where coincidence loss events are consistent. Resolution estimations were made following the FSC 0.143 criteria (Rosenthal & Henderson, 2003) and results are plotted as a function of percent of physical Nyquist frequency versus particle number. The highest achievable resolution is labelled. Data was processed in RELION v3.1, following standard protocols (Scheres, 2012; Zivanov *et al*., 2018).

### Imaging aldolase with and without CDS mode

As a check on the generality of our apoferritin observations, we also explored the influence of dose rate and CDS mode on aldolase, which is smaller (∼150 kDa) and has less symmetry (D2). Unlike the apoferritin sample, aldolase was used as purchased rather than expressed and purified in-house, a fact that potentially limits the ultimate achievable resolution. Consistent with the apoferritin data sets, collecting at 8eps outperformed 16eps in terms of achievable Nyquist frequency. Lower dose rate yielded higher resolution reconstructions across particle subsets, with a larger difference being seen when fewer particles were used. Reconstructions from 10,000-particle subsets reached 3.5 Å (77%) at 8eps vs. 3.8 Å (71%) at 16eps (Figure S6 purple and orange, Table S8).

Surprisingly, the gain from using CDS mode when coincidence loss was constant was trivial compared with apoferritin. Reconstructions from data collected at 8eps with CDS yielded nearly the same resolutions as 16eps without CDS across the different subsets (Figure S6 purple and yellow curves, Table S8). By contrast, at fixed dose rates, CDS was always comparable or better than without CDS (Figure S6 purple vs pink or red vs orange, Table S8). For data collected at 8eps, CDS mode is 5-10% better than nonCDS with the 200,000 particle subsets reconstructing to 2.7 Å (CDS) and 2.8 Å (nonCDS), respectively. Thus, the overall trends seen for apoferritin held for aldolase although some confounding effects can be seen presumably due to differences in how SNR vs resolution affect alignment accuracy under the different conditions.

### Trade-off between data collection time and resolution

We have demonstrated that using 9-hole image shift data collection, one can approach (105kx) or significantly surpass (64kx) physical Nyquist frequency using the K3 in CDS mode. Among all settings explored, 8eps CDS consistently yielded the highest quality reconstructions (Figures 1 and 3). While some biological questions necessitate solving the highest resolution structure possible, others benefit from determining multiple structures quickly, or require very large data sets to resolve conformational heterogeneity. We wanted to address the practical question of the trade-offs of collecting data by imaging more quickly at a higher dose rate vs collecting more particles per micrograph at a larger pixel size. We analysed data collection time versus the achievable resolution by processing subsets of micrographs using a quick, standard cryoSPARC data processing pipeline and comparing the reconstruction resolution versus active data collection time (Figure 4).

**Figure 4.**
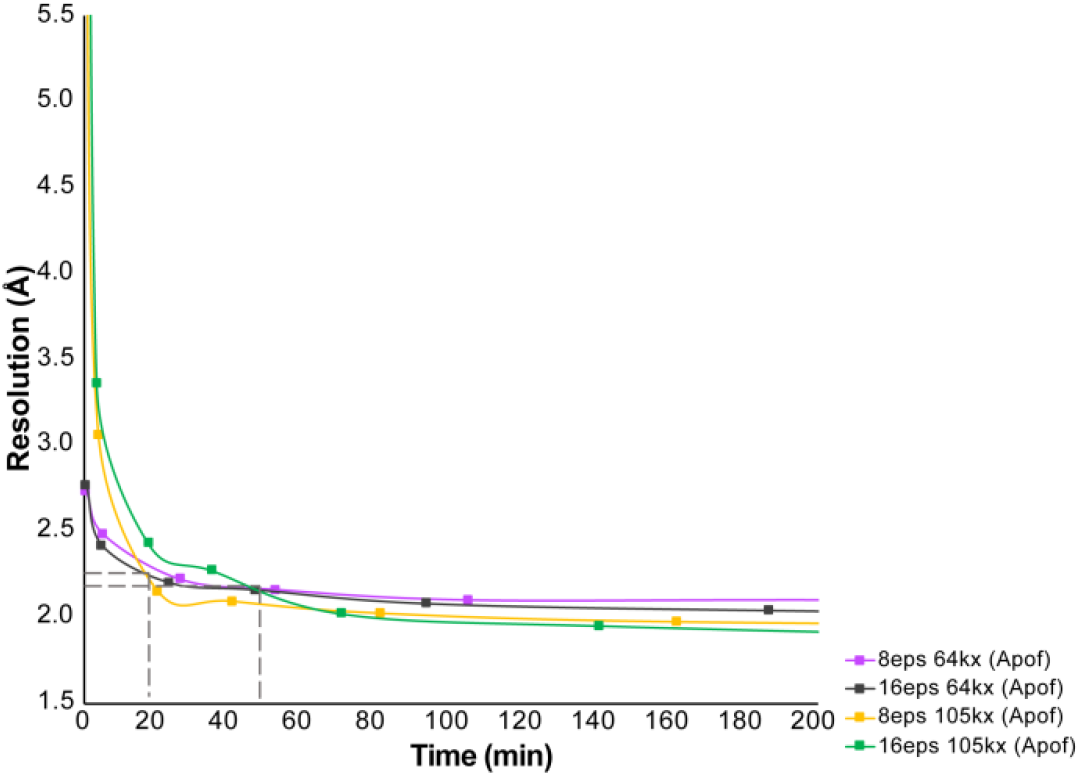
Time versus resolution plot for apoferritin refined with octahedral symmetry. 35% of particles were selected from each image to determine the time it takes different data collection conditions to reach a certain resolution. There is a crossover point where a smaller pixel size should be utilized to reach the highest resolutions, <2.2 Å resolution, between 15- and 50-min depending on dose rate. Data was processed in cryoSPARC v2.12.4 4 (Punjani *et al*., 2017), following standard protocols without 3D classification.

For apoferritin, five minutes of data collection yielded the necessary number of particles to reconstruct below 3 Å (Figure 4). Interestingly, comparing the two magnifications a crossover point was observed where it became more advantageous to image apoferritin at high magnification. This crossover point was at ∼2.2 Å and between 20-60 minutes of data collection time. The data indicate that imaging at a 64kx magnification reaches 2.5 Å resolution ∼5x faster than imaging at 105kx magnification. Thus, an intermediate resolution data collection session could last as little as 30 minutes at 64kx magnification, not including automated data collection set-up, for this benchmark sample. To reach high resolutions, below 2 Å for apoferritin, a higher magnification should be used and the data collection session will need to be at least 2.5 hours (Figure 4). If octahedral symmetry is not applied, making the sample more comparable to other large but asymmetric particles, a data collection of around 8 hours would be necessary to reach sub-2 Å resolution (Figure S3).

### Maximal resolutions attainable with these data and atomic modelling

To eliminate the effects of unexplored higher-order optical aberrations and astigmatisms that were present in the dataset, we performed three rounds of CTF refinements for all the apoferritin and aldolase data sets: refining magnification anisotropy, then refining third and fourth-order optical aberrations, and finally refining per-particle defocus and per-micrograph astigmatism. With these, we far surpassed the physical Nyquist limit of 2.69 Å for apoferritin data collected at 64kx magnification (1.9 Å) and reached 1.75 Å, 95% of the physical Nyquist limit of 1.69Å at 105kx magnification, using 200,000 particles.

To further optimize motion correction for the data sets collected at the optimal dose rate, 8eps, we performed Bayesian polishing, followed by 3D auto refinement (Figure S4 and Table S7). These steps improved the overall resolution of the reconstructions collected at all magnifications and in both CDS and nonCDS mode, as expected. It is worth noting that for the highest resolution reconstructions from 200k and 571k particles at 105kx magnification, polishing improved the resolution to 1.69 Å and 1.65 Å, respectively. Similarly, polishing improved the resolution at 64kx magnification from 1.91Å to 1.87Å (200k particles) and from 1.86 Å to 1.80Å (373k particles). The latter corresponds to 149% of the physical Nyquist resolution, indicating the remarkable benefits of super-resolution in CDS mode.

For fitting the atomic model at this resolution, we adopted high-resolution refinement strategies from traditional x-ray crystallography and performed such refinements with individual b-factor refinements in reciprocal space. In such, we were able to fit each atom, including hydrogens, with appropriate per-atom B-factors. When viewing our highest resolution map, density for hydrogen atoms is clearly visible on both the alpha carbon backbone and some side chains (Figure 5D, E, I, J). They are better resolved at 1.65 Å as expected. In fact, some hydrogens are more visible in our cryo-EM maps than maps simulated at 1.5 Å and 1.3 Å resolution would predict (Figure 5A-C, F-H).

**Figure 5.**
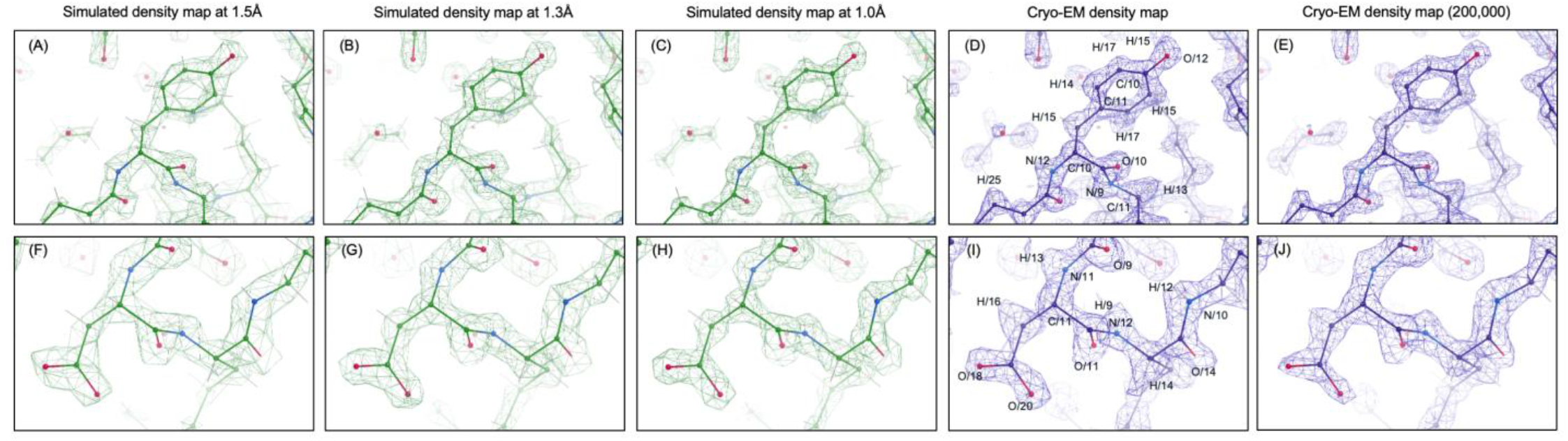
Structural features of apoferritin cryo-EM density map at a global resolution of 1.65 Å. Two apoferritin residues, Trp-168 **(A-E)** and Asp-171 **(F-J)** are selected for comparison among our cryo-EM reconstructions from different number of particles (reconstruction from 500k particles **(D)** and **(I)**; reconstruction from 200k particles **(E)** and **(J)**) and simulated density maps at the indicated resolution **(A-C and F-H)**. The refined atomic coordinates are fitted into each density map and shown as stick and ball. Density that can be attributed to hydrogen atoms, specifically HB2/ Trp 168 (assigned b-factor ∼15) and HD1 Trp 168 (assigned b-factor ∼14) in tops panels and HB2/Asp 171 (assigned b-factor ∼16) and H/ Asp 171 (assigned b-factor ∼13) in bottom panels, can be clearly observed in our cryo-EM density map, in 1.0 Å **(C and G)** and 1.3 Å **(B and F)** simulated maps, but are barely seen in the 1.5 Å simulated map **(A and E)**. Maps simulations were carried out in Phenix (Adams *et al*., 2010) using phenix.fmodel and phenix.mtz2mrc functions, which consider per-atom b factors and generate maps at different resolutions. Estimated per atom b-factors were labelled in 5D and 5I. Cryo-EM data was collected at 105kx, 8eps, CDS mode using 9-hole image shift data collection scheme. Figures were prepared in Coot ((Emsley & Cowtan, 2004)) software.

## Discussion

In this study, we experimentally evaluated Gatan K3 camera performance as a function of dose rate, magnification and CDS mode using the standard sample, apoferritin. In CDS mode, imaging at 8 electron per pixel per second (eps) was always better or as good as imaging at 16eps (Figure 1A and Figure S3), which was expected based on coincidence loss effects. While, surprisingly, reducing the dose rate to 4 eps provided no additional benefit and instead slightly worsened obtainable resolutions. With increasing dose rate, the DQE will decrease, especially at low frequencies. This is largely due to a depressed NPS and MTF whereas higher resolutions are largely unaffected. Comparing reconstructions collected at 8eps and 16eps, the differences in resolutions were greater at the 105kx magnification, than at the 64kx magnification. This suggests that increased coincidence loss affects final reconstruction resolutions most by down weighting intermediate frequencies that are well within physical Nyquist (Li, Zheng *et al*., 2013). By contrast with coincidence loss, the expectation is that CDS mode would enhance DQE across all resolution ranges. Influences of CDS mode at different coincidence loss rates were characterized by comparing reconstructions from apoferritin data sets collected at the same coincidence loss rate or same read-out dose rate (2x difference in terms of coincidence loss rate). Our results validated that operating in CDS mode would further reduce noise and improve DQE and in such, compensates for any increased coincidence losses, in general (Figure 2 and Figure 3).

There have been many discussions as how close obtainable resolutions can be to those imposed by the physical Nyquist sampling. Of concern are both aliasing artifacts, where higher resolution information becomes folded into lower resolution observations, as well as optimal utilization of the camera DQE that falls off as physical Nyquist is approached. The advent of super-resolution sub-pixel localization on the K2 camera provided a straightforward approach to avoiding aliasing. However, the DQE at physical Nyquist remains low, significantly diminishing information transfer beyond this point. Indeed, most efforts to obtain high resolution reconstructions of apoferritin have used physical pixel sizes of 0.46 Å or smaller (Nakane *et al*., 2020; Yip *et al*., 2020; Zhang *et al*., 2020).

Fortunately, the many improvements in the K3 camera significantly increase DQE performance at Nyquist frequency, forcing us to reconsider the true utility of super-resolution information. This is clearly evident in the apoferritin data collected at 64kx magnification. Using 8eps and CDS mode, we readily obtained 1.91 Å or 141% of the physical Nyquist limit (2.69 Å) with 200k particles. Using a smaller subset, of 30k particles, reached 132% of Nyquist, whereas the full set of good particles (373k) combined with Bayesian polishing resulted in a final resolution of 1.80 Å, corresponding to a stunning 149% of the physical Nyquist. This indicates that the increased DQE now available in the K3, especially when using CDS mode, can significantly benefit data collected at lower magnification than typically utilized for high-resolution cryo-EM.

A parallel study on aldolase validated the benefit of using CDS mode for high-resolution cryo-EM reconstruction determination and are largely consistent with apoferritin with a few notable differences. As expected, collecting data at a higher dose rate consistently worsened achievable resolution, but the negative effects of increasing dose rates were larger than observed with apoferritin. (Figure S6). When dose rates were matched, CDS clearly outperformed nonCDS mode for both apoferritin (Figure 3) and aldolase (Figure S6). However, when the coincidence loss rate remained constant, CDS only had a trivial impact on the aldolase and seemed unable to overcome the loss from increased coincidence events as it does for apoferritin. This is likely because coincidence losses result in an amplitude decrease at resolutions lower than half of the physical Nyquist limit, and such resolution may dominate the particle alignment accuracy for smaller particles like aldolase.

As to the goal of balancing collection time and resolution, our data showed that imaging at a larger pixel size allowed us to reach intermediate resolutions ∼5x faster than imaging at a higher magnification (Figure 4). This holds true in our imaging scheme because we acquire one image per hole. The benefit of more particles per image would be offset when taking multiple images per hole (Weis & Hagen, 2020). Because reaching intermediate resolutions appears faster at lower magnification, routine collection at a lower magnification would be most desirable when large numbers of particles are required, such as when determining multiple structures of heterogenous samples. By contrast, if the hypothesis being tested can only be addressed with a very high-resolution structure, such as locating the binding site and position of a small molecule or resolving disease mutations traced to single amino acids, high magnification and physical pixel sizes below 1 Å would be preferred.

In summary, high (physical pixel below 1 Å) or low magnification (physical pixel above 1 Å) should be selected based on the biological problem being asked, noting that reconstructions surpassing the physical Nyquist are now readily achievable when using a larger pixel size. At either magnification, we would advise users to routinely use an intermediate dose rate of 8eps in CDS mode and a 9-hole image shift data collection method to achieve both high-resolution reconstructions and high-throughput collections.

## Materials and Methods

### Plasmid and constructs of heavy chain mouse apoferritin

Plasmid: pET24a-mFth1 (full-length mouse Ferritin heavy chain w/o tag): AAGTGGCGAGCCCGATCTTCCCCATCGGTGATGTCGGCGATATAGGCGCCAGCAACCGCACCTGTGGCGCCGGTGATGCCGGCCACGATGCGTCCGGCGTAGAGGATCGAGATCTCGATCCCGCGAAATTAATACGACTCACTATAGGGGAATTGTGAGCGGATAACAATTCCCCTCTAGAAATAATTTTGTTTAACTTTAAGAAGGAGATATACATATGACCACCGCGTCTCCCTCGCAAGTGCGCCAGAACTACCACCAGGACGCGGAGGCTGCCATCAACCGCCAGATCAACCTGGAGTTGTATGCCTCCTACGTCTATCTGTCTATGTCTTGTTATTTTGACCGAGATGATGTGGCTCTGAAGAACTTTGCCAAATACTTTCTCCACCAATCTCATGAGGAGAGGGAGCATGCCGAGAAACTGATGAAGCTGCAGAACCAGCGAGGTGGCCGAATCTTCCTGCAGGATATAAAGAAACCAGACCGTGATGACTGGGAGAGCGGGCTGAATGCAATGGAGTGTGCACTGCACTTGGAAAAGAGTGTGAATCAGTCACTACTGGAACTGCACAAACTGGCTACTGACAAGAATGATCCCCACTTATGTGACTTCATTGAGACGTATTATCTGAGTGAACAGGTGAAATCCATTAAAGAACTGGGTGACCACGTGACCAACTTACGCAAGATGGGTGCCCCTGAAGCTGGCATGGCAGAATATCTCTTTGACAAGCACACCCTGGGACACGGTGATGAGAGCTAACTCGAGCACCACCACCACCACCACTGAGATCCGGCTGCTAACAAAGCCCGAAAGGAAGCTGAGTTGGCTGCTGCCACCGCTGAGCAATAACTAGCATAACCCCTTGGGGCCTCTAAACGGGTCTTGAGGGGTTTTTTGCTGAAAGGAGGAACTATATCCG GAT

### Protein Expression and Purification

#### Heavy chain mouse apoferritin

Murine heavy chain ferritin in pET24a plasmid was gifted (M. Kikkawa), amplified in DH5a, and sequenced (ElimBio). One liter of *E. coli* BL21(DE3) was grown in LB with 50 ug/mL kanamycin and 0.5% glucose to OD_600_ 0.75. 1 mM IPTG was added for 2.5 hr at 37°C. Cells were harvested, washed in phosphate buffered saline, and resuspended in 30 mM HEPES pH 7.5, 300 mM NaCl, 1 mM MgSO4, 1mg/mL lysozyme, and an EDTA-free protease inhibitor tab (Sigma). The resuspension was sonicated for 150 seconds at 50% power (120-watt Active Motif). Cell debris was centrifuged at 20,000 x*g* for 30 minutes at 4°C. The supernatant was incubated at 70°C with gentle nutation. The sample was centrifuged at 20,000 x*g* for 30 minutes at 4°C. 6.65 g of (NH_4_)_2_SO_4_ was added to 20 mL of the supernatant on ice and stirred for 10 minutes. The sample was centrifuged at 14,000 x*g* for 20 minutes and the pellet was resuspended in 2 mL 20 mM HEPES pH 7.5, 300 mM NaCl. The sample was then dialyzed against 20 mM HEPES pH 7.5, 300 mM NaCl overnight at 4°C in a 20,000 MWCO Slide-A-Lyzer (ThermoFisher). The sample was recovered from the dialysis cassette and centrifuged at 55,000 x*g* for 15 minutes at 4°C. The supernatant was concentrated to ∼1 mL in a 50 MWCO spin column (Millipore). The final concentrated sample was centrifuged at 55,000 x*g* for 15 minutes at 4°C. An A_280_ reading was taken in order to not overload the Superose 6/150 Increase column. The sample was run over a the Superose 6/150 Increase column in two aliquots at 0.4 ml/min in 20 mM HEPES pH 7.5, 300 mM NaCl. Peak fractions were saved and frozen at - 80°C in aliquots at 13 mg/mL (first run) and 6 mg/mL (second run) as determined by nanodrop A_280_ readings.

#### Rabbit aldolase

Rabbit aldolase was purchased from Sigma-Aldrich (A2714-100UN, lot# SLBR7752V) in powdered form. Sample was dissolved at 1.6 mg/mL in 20 mM HEPES (pH 7.5), 50 mM NaCl and used directly.

### Cryo-EM Grid Preparation

To closely mimic standard sample preparation practices in the facility, all apoferritin and aldolase samples were frozen on 300-mesh Quantifoil gold holey carbon grids with hole size and spacing of 1.2/1.3 μm (Quantifoil) using a Mark IV vitrobot (ThermoFisher). Freezing details described as the followings.

#### Apoferritin

6 mg/mL apoferritin aliquots were thawed from storage at -80°C and centrifuged at 55,000 x*g* for 15 minutes at 4°C. 2 μl apoferritin was applied to glow-discharged gold holey carbon 1.2/1.3 300-mesh grids (Quantifoil). Grids were blotted for 4 seconds at -2 force and vitrified in liquid ethane using a MarkIV Vitrobot (ThermoFisher). The blotting chamber was maintained at 22°C and 100% humidity during freezing.

#### Aldolase

0.8 mg/mL aldolase solution was applied to glow-discharged gold holey carbon 1.2/1.3 200 mesh grids (Quantifoil). Grids were blotted for 3 seconds at 0 force and vitrified in liquid ethane using a MarkIV Vitrobot (ThermoFisher). The blotting chamber was maintained at 6°C and 100% humidity during freezing.

### Cryo-EM Data Collection

Briefly, all images were acquired on a Titan Krios microscope outfitted with a Gatan K3 detector and BioQuantum Imaging Filter operating in nanoprobe and EF-TEM mode, a C2 aperture size of 70 µm, an objective aperture size of 100 µm, and an energy filter slit width of 20 eV. Data was collected automatically using 3×3 9-hole image shift application in SerialEM, with the same defocus range, total electron dose, and frame number with a constant electron dose per frame. Details are described as the followings.

#### Microscope alignment

Full alignments of the microscope were performed only after K3 installation, after which only direct alignments were routinely checked before each data collection. On a Titan Krios microscope, using a 70 µm C2 aperture, the sample was first set to a stage position at eucentric height over carbon on a grid square that appeared free of defects at lower magnification. Specimen was then brought close to focus and astigmatism was adjusted before proceeding with beam alignment. First, pivot points and rotation center were roughly adjusted at imaging magnification (or higher at nominal 165,000x for rotation center). Further tuning of pivot points and rotation center were then performed by iterating through microscope coma-free pivot point and coma-free alignment procedures for four directions of beam tilt. Finally, SerialEM implementation of astigmatism and Coma-free alignment correction by CTF fitting was used for fine adjustment, correcting 8 beam tilt directions. Beam intensity was adjusted to a diameter within parallel illumination conditions for a 3-condenser lens system, and gave the appropriate dose for the dataset (spot size varying between 4 and 5). SerialEM Coma vs. Image Shift procedure was then performed for each of these dose settings accounting for any differences in illumination and saved as separate settings files.

#### BioQuantum Gatan energy filter tuning

The BioQuantum Gatan energy filter was tuned once a week, without sample in the beam path. We found that within a week for our GIF values for isochromaticity, achromaticity, magnification and distortion had almost no change (within 1 eV for isochromaticity). tuning was performed by decreasing spot size to 1 which would give sufficient dose for full GIF tuning to yield good results and uniform slit edge. For zero-loss peak centering, a dose around 16 eps was sufficient and consistent within 5eV over the course of a week. A slit width of 20 eV was used for data collection. A 100 μm objective aperture was then inserted and centered before a final astigmatism adjustment, after which the microscope was ready for data collection.

#### Considerations of data collection consistency

For consistency, apoferritin datasets were collected in one imaging session, over the course of four days, on a single grid. Aldolase datasets were collected in two sessions from grids vitrified under the same conditions in the same freezing session. A gain reference was collected in CDS and nonCDS mode for each magnification and dose rate.

### Image processing

During data collection, images were pre-processed to provide feedback on image quality. All pre-processing was carried out using Scipion software (de la Rosa-TrevÍn *et al*., 2016). Beam-induced motion correction and dose weighing were performed on the raw movie stacks using MotionCor2 (Zheng *et al*., 2017) using a 7×5 patch size and a b-factor of 100 with 7 iterations. Super-resolution movies were saved for further analysis. Whole-image contrast transfer function (CTF) estimation was performed using CTFFFIND 4 (Rohou & Grigorieff, 2015).

#### Apoferritin data collected at 105,000x magnification

For the data sets of apoferritin, collected at **105**,**000 x magnification, dose rates of 4 and 8 and 16 eps**, image processing was performed in RELION 3.1 (Zivanov *et al*., 2018; Scheres, 2012) software following the standard protocols. Particles were picked using the Laplacian-of-gaussian algorithm and extracted at 4x binned. These particles were subjected to standard 3D classification, to eliminate any unfit classes. The selected good class was re-extracted at 1x binned with a box size of 512 pixels and subjected to standard 3D auto-refinement to give the initial reconstruction. Subsequently, three runs of CTF refinement were performed using default parameters: first refining magnification anisotropy; then refining optical aberrations (up to the 4th order); and finally refining per-particle defocus and per-micrograph astigmatism. Another round of 3D auto-refinement with the refined values were performed.

#### High-resolution structure of apoferritin

To further improve the resolution of the optimal data set, the one collected with a dose rate of 8eps at 105,000x magnification at CDS mode, the standard Bayesian polishing (Zivanov *et al*., 2019) was performed, followed by standard 3D auto-refinement. Then another round of CTF refinement was performed as it’s described earlier. These yield a final average resolution of 1.66 Å (FSC=0.143), with an estimated accuracy aligned angles 0.325 degrees and offsets= 0.182969 Angstroms.

#### Apoferritin data collected at 64,000x magnification

For the data sets of **apoferritin** collected at **64**,**000x magnification, dose rates of 8 eps under CDS and standard nonCDS modes**, particles were picked using Gautomatch (Kai Zhang, https://www.mrc-lmb.cam.ac.uk/kzhang/). The following image processing was performed in RELION 3.1 (Zivanov *et al*., 2018; Scheres, 2012) software using the same protocols described above. For the data sets of **apoferritin** collected at **64**,**000 magnification, dose rates of 16 eps under CDS and standard non-CDS modes, image processing was performed** using cryoSPARC 2.14.2 (Punjani *et al*., 2017) following standard protocols. Particles were picked using the template picking algorithm using preliminary 2D averages from a blob pick job performed on 10 micrographs. Putative particles were extracted and 8x binned. 2D classification was performed to screen out ice contamination and clearly damaged particles. The remaining particles were subjected to Ab Initio Reconstruction with octahedral symmetry, to eliminate any unfit classes. The good class was re-extracted without binning at a box size of 300 pixels and subjected to standard Homogenous Refinement including Defocus Refinement and Global CTF Refinement using octahedral symmetry. All available higher-order magnification parameters were fit and corrected.

Final particle stack was exported using csparc2star.py (Asarnow, D. et al., 2019), masked, and postprocessed in RELION 3.1 (Zivanov et al., 2018; Scheres, 2012).

All reported resolutions were based on the 0.143 Fourier shell criterion (Chen *et al*., 2013; Rosenthal & Henderson, 2003) with all Fourier shell correlation (FSC) curves corrected for the effects of soft solvent masking by high-resolution noise substitution (Chen *et al*., 2013) as calculated in RELION 3.1. Masks were generated in RELION at the 50% of the density maps’ maximum density values with 8-pixel extension and 8-pixel width.

#### Aldolase data collected at 64,000x magnification

For the data sets of **aldolases**, collected at **64**,**000 x magnification, dose rates of 8eps under CDS and standard nonCDS modes**, image processing was performed in RELION 3.1 software following the same standard as for apoferritin data collected at 64,000x magnification. For the data sets of aldolases collected at **64**,**000x magnification, dose rates of 16eps under CDS and standard non-CDS modes**, image processing was performed using cryoSPARC 2.14.2 (Punjani *et al*., 2017) following the same standard as for apoferritin collected at 64,000x magnification.

All reported resolutions are based on the 0.143 Fourier shell criterion (Chen *et al*., 2013; Rosenthal & Henderson, 2003) with all Fourier shell correlation (FSC) curves corrected for the effects of soft solvent masking by high-resolution noise substitution (Chen *et al*., 2013). Masks were auto generated in cryoSPARC and postprocessing was run in RELION for consistency with data sets processed in RELION.

#### Time to resolution plots

Processing was carried out as above except particles were not subjected to a 3D classification step using Ab Initio Reconstruction job and per-particle defocus and local CTF refinement were not performed. These particles were selected, cleaned from obvious contamination and refined using standard Homogenous Refinement in cryoSPARC (Punjani *et al*., 2017) to achieve a reasonable resolution in the shortest period of time. For apoferritin refinements with octahedral symmetry, ∼35% of particles were selected using the Inspect Picks job in cryoSPARC because when using all particles in an image the resolution dropped too quickly to see any difference between different conditions. For apoferritin refinements with C1 symmetry and all aldolase refinements, all picked particles were used. Final resolutions are reported from the tight mask FSC curves reported from cryoSPARC 2.14.2 (Punjani *et al*., 2017).

### Atomic modelling

The x-ray crystal structure PDB 5OBA was docked into the cryo-EM density map in UCSF Chimera (Pettersen *et al*., 2004) and the 24 chains were refined against full the cryo-EM density maps in Phenix software (Adams *et al*., 2010) following the recommended protocol for refining medium to high resolution structures. In reciprocal space refinement, the main refinement loops are as followings, (1) converting cryo-EM density map to structure factors using phenix.map_to_structure_factors function; (2) refinement with bulk-solvent and individual anisotropic refinement using phenix.refine function; (3) refinements with individual anisotropic refinement but without bulk-solvent refinement using phenix.refine function. In addition, during the individual anisotropic refinement, the hydrogens were refined as a riding model. The model was then further refined in real space with phenix.real_space_refine and subsequently adjusted manually in Coot (Emsley & Cowtan, 2004). Validation was carried out using full validation function in Phenix. The full cryo-EM density map with intermediate b-factor sharpened applied, b factor = -25, was used for phenix.map_to_structure_factors conversion and cryo-EM validation.

### Calibration and refinement of pixel sizes

Gatan K3 pixel size was initially calibrated through an autocorrelation routine in SerialEM (Mastronarde, 2005) using a cross-line replica grating grid, finding correlation peaks progressively further out to estimate the spacing between peaks along two axes. Using this method, we estimated the pixel size value at both nominal magnifications of 105,000 x and 64,000 x to be 0.853 Å/pixel and 1.378 Å/pixel initially. We then used the high-resolution reconstructions of apoferritin to further refine the pixel size (Wu et al., 2020). The x-ray crystal structures of apoferritin were first docked and then refined into respective cryo-EM reconstructions of voxel sizes ranging from 0.416 Å/pixel (counted 0.832 Å/pixel) to 0.4265 Å/pixel (counted 0.853 Å/pixel) for data collected at 105kx magnification, and from 0.671 Å/pixel (counted 1.342 Å/pixel) to 0.689 Å/pixel (counted 1.378Å/pixel) using phenix.real_space_refinement package (Adams et al., 2010). Meanwhile, the x-ray structure was also rigid-body fitted into the cryo-EM density maps in UCSF Chimera (Pettersen et al., 2004) By examining the map vs map CC values reported from phenix and the x-ray simulated map vs cryo-EM CC values from Chimera, we found that at the pixel size values of 0.834 Å/pixel and 1.346 Å/pixel gave the highest score.

## Author Contributions and Notes

M.S, C.M.A, E.T, D.A.A, K.A.V, Y.C, D.S, and A.F designed research, M.S, C.M.A, and E.T performed research, M.S, C.M.A, and E.T analyzed data; and M.S, C.M.A, E.T, D.A.A, K.A.V, Y.C, D.S, and A.F wrote and edited the paper.

The authors declare no conflict of interest.

## Data availability

The electron density maps and atomic models have been deposited into the Electron Microscopy Data Bank (EMDB) and the Protein Data Bank (PDB). The accession codes are EMD-22972 and 7KOD, respectively.

## Acknowledgments

We thank the Kikkawa lab for the murine heavy-chain ferritin construct plasmid; David Bulkley and Glenn Gilbert for support in data collection and microscope management through the University of California San Francisco Advanced Cryo-Electron Microscopy Facility; Matt Harrington, Joshua Baker-LePain, and the QB3 shared cluster (NIH grant 1S10OD021596-01) for high-performance computing (HPC) support; Paul Mooney and Chris Booth (Ametek Gatan) for insightful discussions. The software used for this project was curated by SBGrid (Morin et al., 2013).

This work was supported by the National Institutes of Health, National Institute of General Medical Sciences (grant No. R35GM118099 to David A. Agard; grant No. R01HL134183 to David A. Agard; grant No. P50AI150476 to Yifan Cheng; grant No. S10OD020054 to UCSF Cryo-EM facility; grant No. S10OD021741 to UCSF Cryo-EM facility; grant No. S10OD026881 to UCSF Cryo-EM facility). Y.C. is an Investigator of Howard Hughes Medical Institute. A.F. is supported in part by a faculty scholar grant from the Howard Hughes medical institute and a Chan Zuckerberg Biohub investigator award.

## Supplementary Information

**Table S1.**
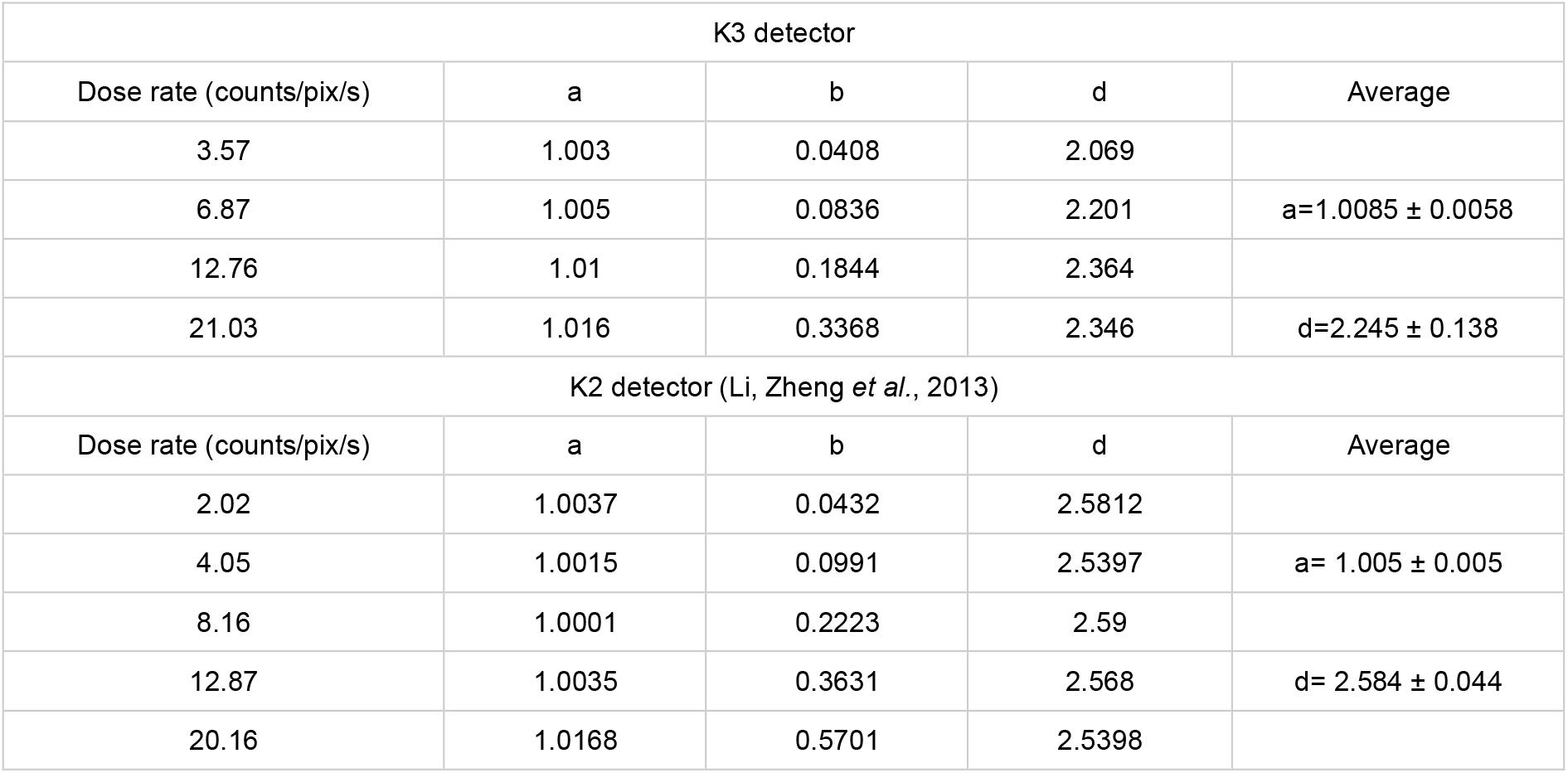
Summary of NPS curve fitting (Relevant to Figure S2). Least squares fitting of the NPS curves obtained from the K3 detector using the function: Intensity = a - b * sinc2(frequency * d) (Li, Zheng *et al*., 2013) equation (1)). Results obtained previously from K2 are shown for comparison. The dampening effect of b reflects coincidence loss and is improved in the K3.

| K3 detector |  |  |  |  |
| --- | --- | --- | --- | --- |
| Dose rate (counts/pix/s) | a | b | d | Average |
| 3.57 | 1.003 | 0.0408 | 2.069 |  |
| 6.87 | 1.005 | 0.0836 | 2.201 | a=1.0085 ± 0.0058 |
| 12.76 | 1.01 | 0.1844 | 2.364 |  |
| 21.03 | 1.016 | 0.3368 | 2.346 | d=2.245 ± 0.138 |
| K2 detector (Li, Zheng *et al.*, 2013) |  |  |  |  |
| Dose rate (counts/pix/s) | a | b | d | Average |
| 2.02 | 1.0037 | 0.0432 | 2.5812 |  |
| 4.05 | 1.0015 | 0.0991 | 2.5397 | a= 1.005 ± 0.005 |
| 8.16 | 1.0001 | 0.2223 | 2.59 |  |
| 12.87 | 1.0035 | 0.3631 | 2.568 | d= 2.584 ± 0.044 |
| 20.16 | 1.0168 | 0.5701 | 2.5398 |  |

**Table S2.** Probability of number of electron events (k) overlapping for the K3 utilizing the effective internal frame rates with CDS mode off or on (Relevant to Table S1). Highlighted are two dose rates with the same probabilities of coincidence loss between the two modes. We used the binomial distribution model described previously (Li, Zheng *et al*., 2013) and the electron ‘mask’ d derived from fitting the intensities of NPS as the area to consider electron overlaps (**Table S1**).

| NonCDS mode (1500 frames per second) |  |  |  |
| --- | --- | --- | --- |
| Dose rate (eps) | k = 0 | 1 | 2 |
| 1 | 99.69% | 0.31% | 0.00% |
| 2 | 99.39% | 0.61% | 0.00% |
| 4 | 98.78% | 1.21% | 0.01% |
| 8 | 97.58% | 2.39% | 0.03% |
| **16** | **95.21%** | **4.67%** | **0.11%** |
| 32 | 90.65% | 8.90% | 0.44% |
| CDS mode (750 frames per second) |  |  |  |
| Dose rate (eps) | k = 0 | 1 | 2 |
| 1 | 99.39% | 0.61% | 0.00% |
| 2 | 98.78% | 1.21% | 0.01% |
| 4 | 97.58% | 2.39% | 0.03% |
| **8** | **95.21%** | **4.67%** | **0.11%** |
| 16 | 90.65% | 8.90% | 0.44% |
| 32 | 82.18% | 16.13% | 1.58% |

**Table S3.** Influence of dose rate on reconstructions of apoferritin from data recorded on the K3 detector (Relevant to Figure 1A). Results of 3D reconstructions from apoferritin data sets collected at different dose rates. All resolutions are reported in absolute angstrom and percentage of physical Nyquist frequency calculated from gold standard FSC estimates with soft mask.

| Magnification | Physical pixel size | Dose rate (e/pixel/sec) | No. of Particles | Resolution | % of Physical Nyquist |
| --- | --- | --- | --- | --- | --- |
| 105,000 | 0.834 Å/pixel | 4 | 10,000 | 2.05 Å | 81% |
|  |  |  | 30,000 | 1.96 Å | 85% |
|  |  |  | 60,000 | 1.89 Å | 88% |
|  |  |  | 200,000 | 1.78 Å | 94% |
| 105,000 | 0.834 Å/pixel | 8 | 10,000 | 1.98 Å | 84% |
|  |  |  | 30,000 | 1.91 Å | 87% |
|  |  |  | 60,000 | 1.84 Å | 91% |
|  |  |  | 200,000 | 1.75 Å | 95% |
| 105,000 | 0.834 Å/pixel | 16 | 10,000 | 2.16 Å | 77% |
|  |  |  | 30,000 | 2.05 Å | 81% |
|  |  |  | 60,000 | 1.94 Å | 86% |
|  |  |  | 200,000 | 1.78 Å | 94% |
| 64,000 | 1.346 Å/pixel | 8 | 10,000 | 2.22 Å | 121% |
|  |  |  | 30,000 | 2.04 Å | 132% |
|  |  |  | 60,000 | 1.98 Å | 136% |
|  |  |  | 200,000 | 1.91 Å | 141% |
| 64,000 | 1.346 Å/pixel | 16 | 10,000 | 2.32 Å | 116% |
|  |  |  | 30,000 | 2.09 Å | 129% |
|  |  |  | 60,000 | 2.00 Å | 135% |
|  |  |  | 200,000 | 1.91 Å | 141% |

**Table S4.** Benefits of imaging apoferritin in CDS mode when comparing data sets with equivalent coincidence loss of electron events (Relevant to Figure 2). Results of 3D reconstructions from apoferritin data sets collected at constant coincidence loss. All resolutions are reported in absolute angstrom and percentage of physical Nyquist frequency calculated from gold standard FSC estimates with soft masks.

| Magnification | Physical pixel size | Dose rate<br>(e/pixel/sec) | Mode | No. of Particles | Resolution | % of Physical<br>Nyquist |
| --- | --- | --- | --- | --- | --- | --- |
| 64,000 | 1.346 Å/pixel | 8 | CDS | 10,000 | 2.22 Å | 121% |
|  |  |  |  | 30,000 | 2.04 Å | 132% |
|  |  |  |  | 60,000 | 1.98 Å | 136% |
|  |  |  |  | 200,000 | 1.91 Å | 141% |
| 64,000 | 1.346 Å/pixel | 16 | NonCDS | 10,000 | 2.32 Å | 116% |
|  |  |  |  | 30,000 | 2.23 Å | 121% |
|  |  |  |  | 60,000 | 2.11 Å | 128% |
|  |  |  |  | 200,000 | 1.98 Å | 136% |
| 105,000 | 0.834 Å/pixel | 4 | CDS | 10,000 | 2.05 Å | 81% |
|  |  |  |  | 30,000 | 1.96 Å | 85% |
|  |  |  |  | 60,000 | 1.89 Å | 88% |
|  |  |  |  | 200,000 | 1.78 Å | 94% |
| 105,000 | 0.834 Å/pixel | 8 | NonCDS | 10,000 | 2.16 Å | 77% |
|  |  |  |  | 30,000 | 2.03 Å | 82% |
|  |  |  |  | 60,000 | 1.98 Å | 84% |
|  |  |  |  | 200,000 | 1.91 Å | 89% |

**Table S5.**
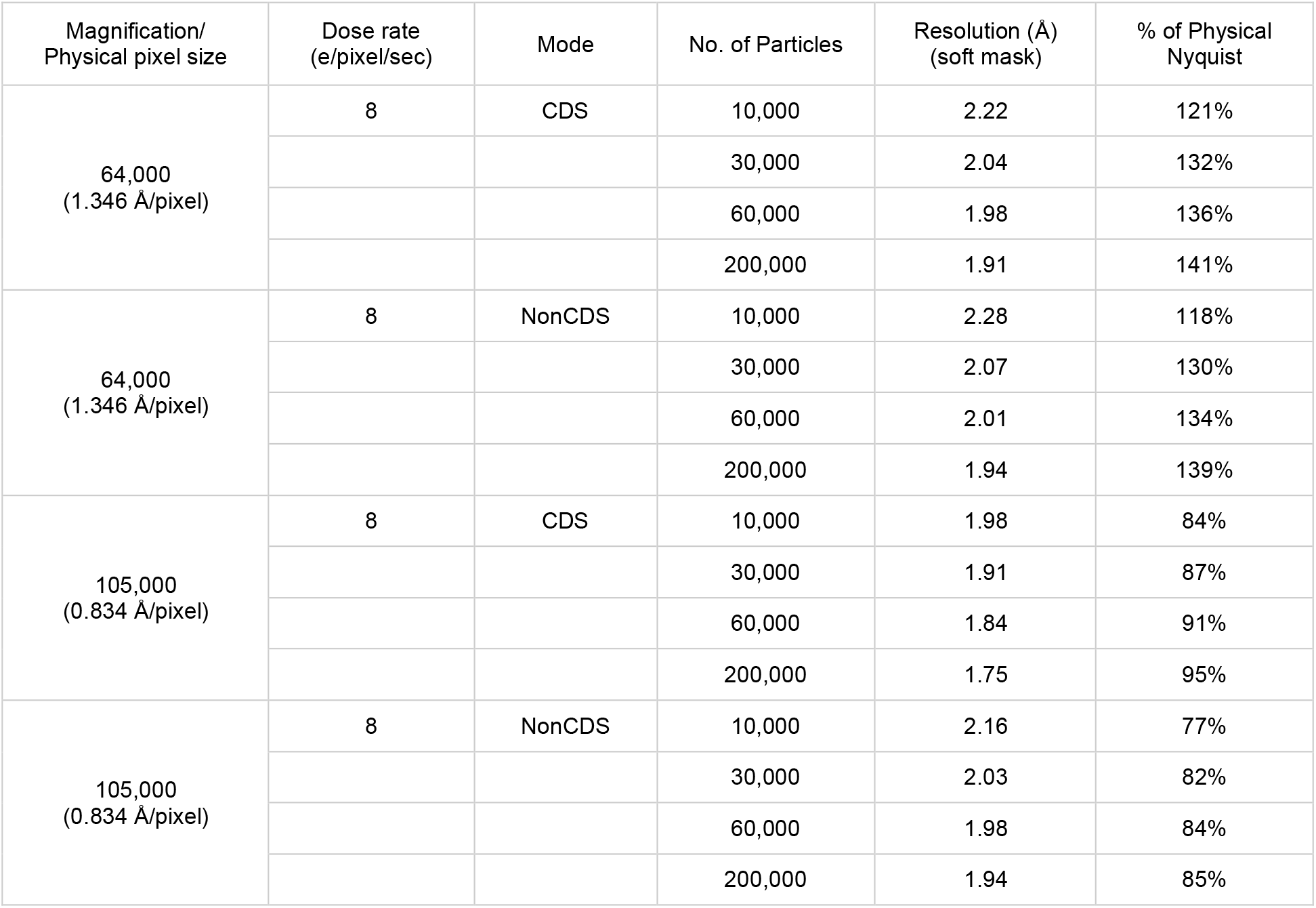
Practical consideration of imaging apoferritin using CDS mode at the recommended dose rate of 8eps (Relevant to Figure 3). Results of 3D reconstructions from apoferritin data sets collected at constant dose rate. All resolutions are reported in absolute angstrom and percentage of physical Nyquist frequency calculated from gold standard FSC estimates (Chen *et al*., 2013; Rosenthal & Henderson, 2003).

**Table S6.** Statistics of data collection, 3D refinement, and atomic model for data collected at 8eps and 105kx in CDS mode (Relevant to Figure 5). The model validation was carried out in Phenix software (Adams *et al*., 2010), using medium sharpened cryo-EM full density map.

| Apoferritin (8eps, 105kx, CDS) |  |
| --- | --- |
| Data collection and processing |  |
| Magnification | 105,000 |
| Voltage (kV) | 300 |
| Electron exposure (e-/Å<sup>2</sup>) | 69 |
| Defocus range (μm) | -0.3 to -0.7 |
| Pixel size (Å) | 0.834 (physical pixel size);<br>0.417 (super) |
| Symmetry imposed | O |
| Map resolution (Å) | 1.65 (FSC 0.143) |
| Refinement |  |
| Initial model used (PDB code) | 5OBA/5OBB |
| Model resolution (Å)<br>(FSC threshold 0.143/0.5) | 1.5/1.7 (0.143/0.5) |
| Model composition<br>Atoms non-H and H<br>Protein residues<br>Water | 68186 (H: 32643)<br>4128<br>1727 |
| B factors (Å<sup>2</sup>) min/max/mean<br>Protein<br>Water | 5/101/21<br>10/66/22 |
| R.m.s. deviations<br>Bond lengths (Å) (#>4σ)<br>Bond angles (°) (#>4σ) | 0.001 (0)<br>0.342 (0) |
| Validation<br>MolProbity score<br>Clashscore<br>Poor rotamers (%) | 0.75<br>0.78<br>0.05 |
| Ramachandran plot<br>Favored (%)<br>Allowed (%)<br>Disallowed (%) | 98.87<br>1.13<br>0.00 |

**Table S7.** Resolution improvements from per-particle motion correction (Relevant to Figure S4). Results of 3D reconstructions from apoferritin and aldolase data sets before and after Bayesian polishing, with and without masking. All resolutions are reported in absolute angstrom and percentage of physical Nyquist frequency calculated from gold standard FSC estimates. All resolutions are reported in absolute angstrom and percentage of physical Nyquist frequency calculated from gold standard FSC estimates (Chen *et al*., 2013; Rosenthal & Henderson, 2003).

| Aldolase 64kx 8eps CDS |  |  |  |  |
| --- | --- | --- | --- | --- |
|  | Auto 3D Refinement |  | Bayesian Polishing |  |
| # of particles | (no mask) | (soft mask) | (no mask) | (soft mask) |
| 252000 | 2.96 Å | 2.66 Å / 99% | 2.88 Å | 2.55 Å / 105% |
| 200000 | 3.01 Å | 2.73 Å / 99% | 2.97 Å | 2.66 Å / 101% |
| Aldolase 64kx 8eps nonCDS |  |  |  |  |
| 252000 | 3.11 Å | 2.69 Å / 100% | 2.92 Å | 2.52 Å / 107% |
| 200000 | 3.15 Å | 2.88 Å / 93% | 2.97 Å | 2.62 Å / 103% |

| Apoferritin 64kx 8eps CDS |  |  |  |  |
| --- | --- | --- | --- | --- |
|  | Auto 3D Refinement |  | Bayesian Polishing |  |
| # of particles | (no mask) | (soft mask) | (no mask) | (soft mask) |
| 373000 | 2.04 Å | 1.86 Å / 144% | 2.01 Å | 1.80 Å / 149% |
| 200000 | 2.11 Å | 1.91 Å / 141% | 2.06 Å | 1.87 Å / 144% |
| Apoferritin 64kx 8eps nonCDS |  |  |  |  |
| 256000 | 2.10 Å | 1.89 Å / 142% | 2.02 Å | 1.87 Å / 144% |
| 200000 | 2.10 Å | 1.94 Å / 139% | 2.02 Å | 1.88 Å / 143% |

**Table S7. Resolution improvements from per-particle motion correction (Relevant to Figure S4).**
| Apoferritin 105kx 8eps CDS |  |  |  |  |
| --- | --- | --- | --- | --- |
|  | Auto 3D Refinement |  | Bayesian Polishing |  |
| # of particles | (no mask) | (soft mask) | (no mask) | (soft mask) |
| 571000 | 1.81 Å | 1.71 Å / 95% | 1.76 Å | 1.65 Å / 101% |
| 200000 | 1.88 Å | 1.75 Å / 95% | 1.79 Å | 1.69 Å / 100% |
| Apoferritin 105kx 8eps nonCDS |  |  |  |  |
| 441000 | 1.97 Å | 1.86 Å | 1.90 Å | 1.76 Å / 95% |
| 200000 | 2.01 Å | 1.87 Å | 1.98 Å | 1.86 Å / 90% |

**Table S8.**
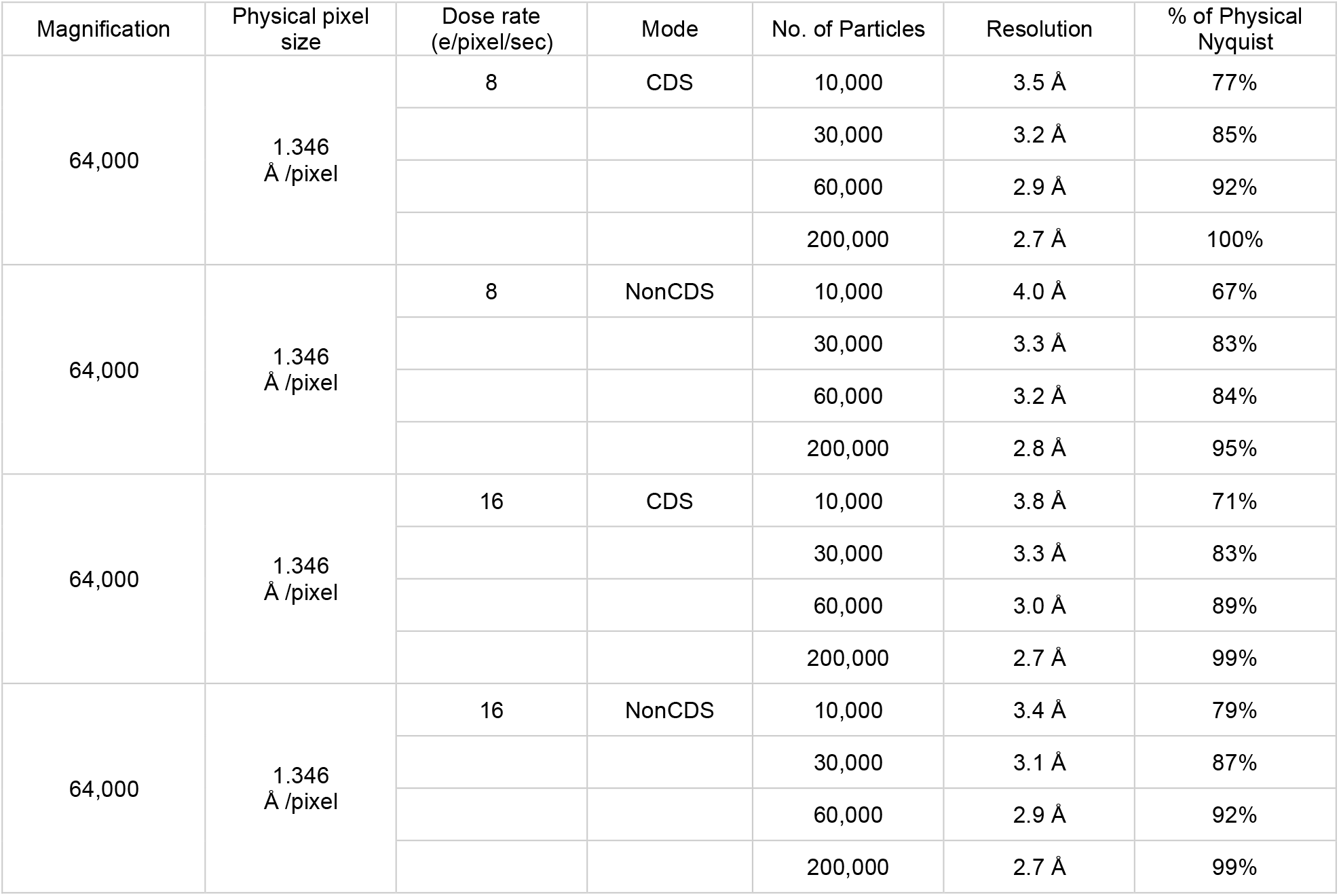
Balancing the effects of dose rate, coincidence loss, and noise reduction when imaging in CDS and nonCDS mode for aldolase (Relevant to Figure S6). Comparison of all aldolase data sets collected at 64,000x magnification. All resolutions are reported in absolute angstrom and percentage of physical Nyquist frequency calculated from gold standard FSC estimates (Chen *et al*., 2013; Rosenthal & Henderson, 2003).

| Magnification | Physical pixel size | Dose rate (e/pixel/sec) | Mode | No. of Particles | Resolution | % of Physical Nyquist |
| --- | --- | --- | --- | --- | --- | --- |
| 64,000 | 1.346 Å /pixel | 8 | CDS | 10,000 | 3.5 Å | 77% |
|  |  |  |  | 30,000 | 3.2 Å | 85% |
|  |  |  |  | 60,000 | 2.9 Å | 92% |
|  |  |  |  | 200,000 | 2.7 Å | 100% |
| 64,000 | 1.346 Å /pixel | 8 | NonCDS | 10,000 | 4.0 Å | 67% |
|  |  |  |  | 30,000 | 3.3 Å | 83% |
|  |  |  |  | 60,000 | 3.2 Å | 84% |
|  |  |  |  | 200,000 | 2.8 Å | 95% |
| 64,000 | 1.346 Å /pixel | 16 | CDS | 10,000 | 3.8 Å | 71% |
|  |  |  |  | 30,000 | 3.3 Å | 83% |
|  |  |  |  | 60,000 | 3.0 Å | 89% |
|  |  |  |  | 200,000 | 2.7 Å | 99% |
| 64,000 | 1.346 Å /pixel | 16 | NonCDS | 10,000 | 3.4 Å | 79% |
|  |  |  |  | 30,000 | 3.1 Å | 87% |
|  |  |  |  | 60,000 | 2.9 Å | 92% |
|  |  |  |  | 200,000 | 2.7 Å | 99% |

**Figure S1.**
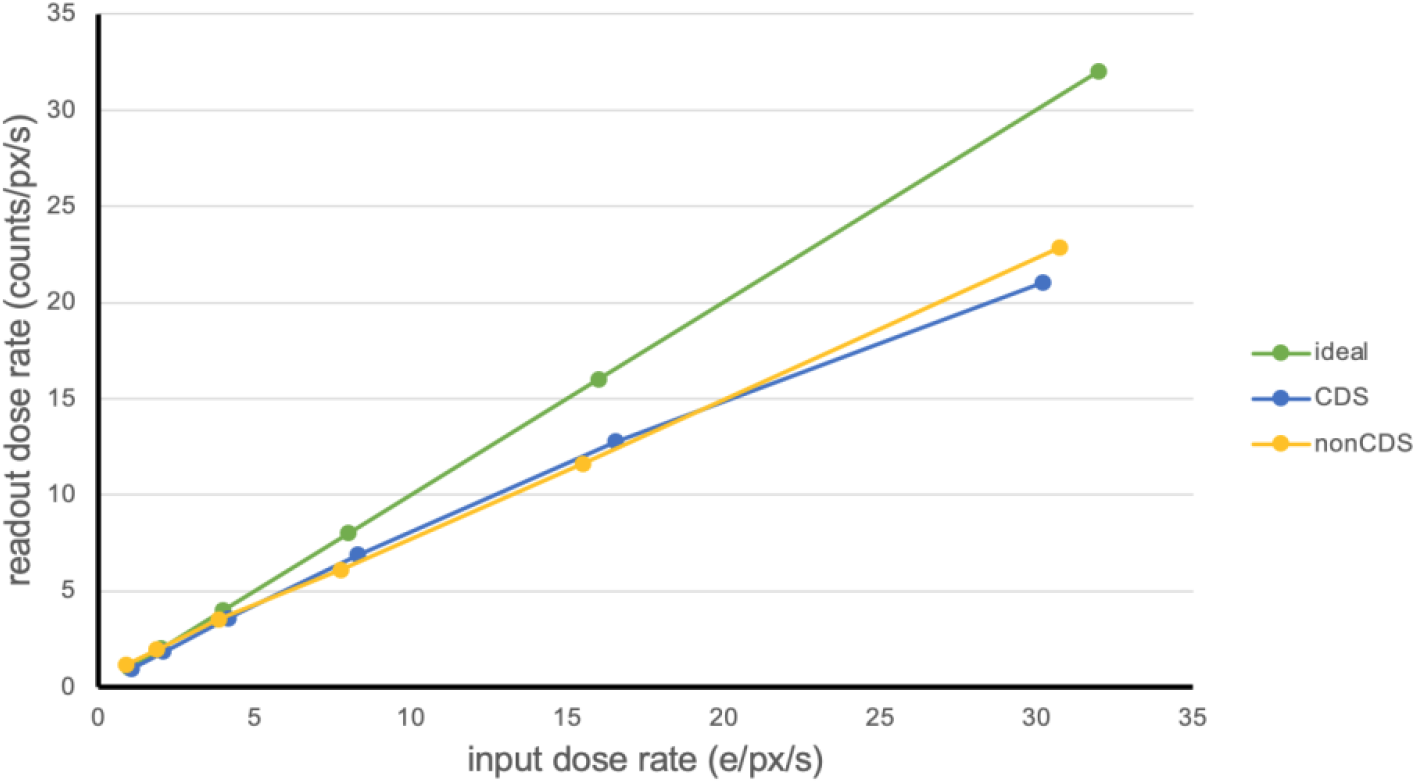
Coincidence loss for the Gatan K3 detector. Detector counts converted to electron dose of the Gatan K3 operating in counted mode with and without CDS mode compared to 1 to 1 conversion. Electron counts can be separated as independent events at lower dose rates which fall off as the incident electron dose rate increases.

**Figure S2.**
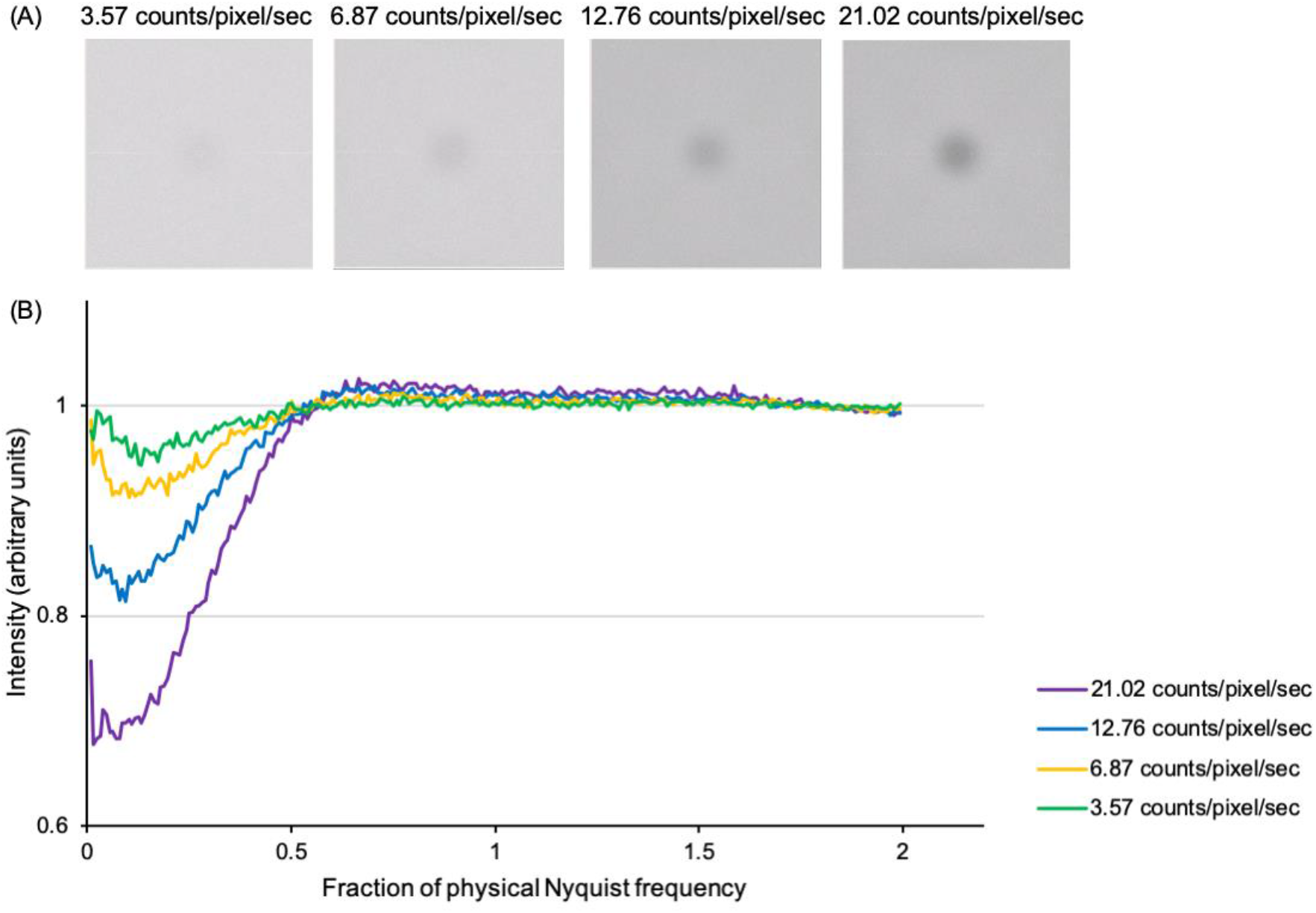
Noise power spectra (NPS) acquired at various dose rates. **(A)** Fourier transforms of empty super-resolution counting images recorded with different dose rates in CDS mode. **(B)** Simulated noise power spectra at different dose rates, by using the normalized sinc function (Li, Zheng *et al*., 2013) equation (1). The dip of the NPS curve at low frequencies describes coincidence loss with increasing dose rate.

**Figure S3.**
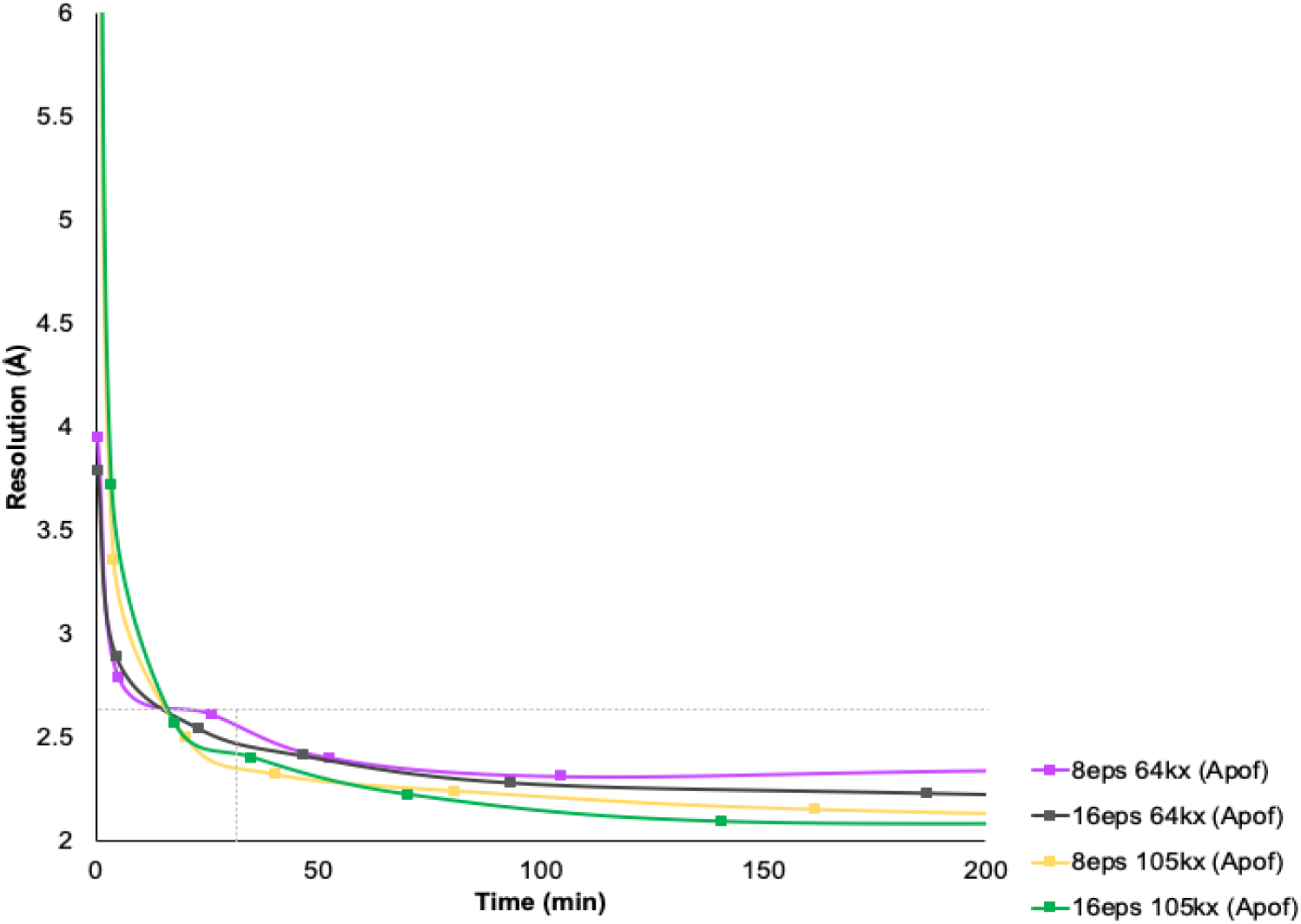
Time versus resolution plot for apoferritin refined with C1 symmetry. All particles from the same selected micrographs in Figure 4 were processed in cryoSPARC v2.12.4 (Punjani *et al*., 2017), following standard protocols without 3D classification as in Figure 4 with C1 symmetry. The 8eps 105kx CDS mode data reconstructed below 2 Å resolution after ∼8 hours of data.

**Figure S4.**
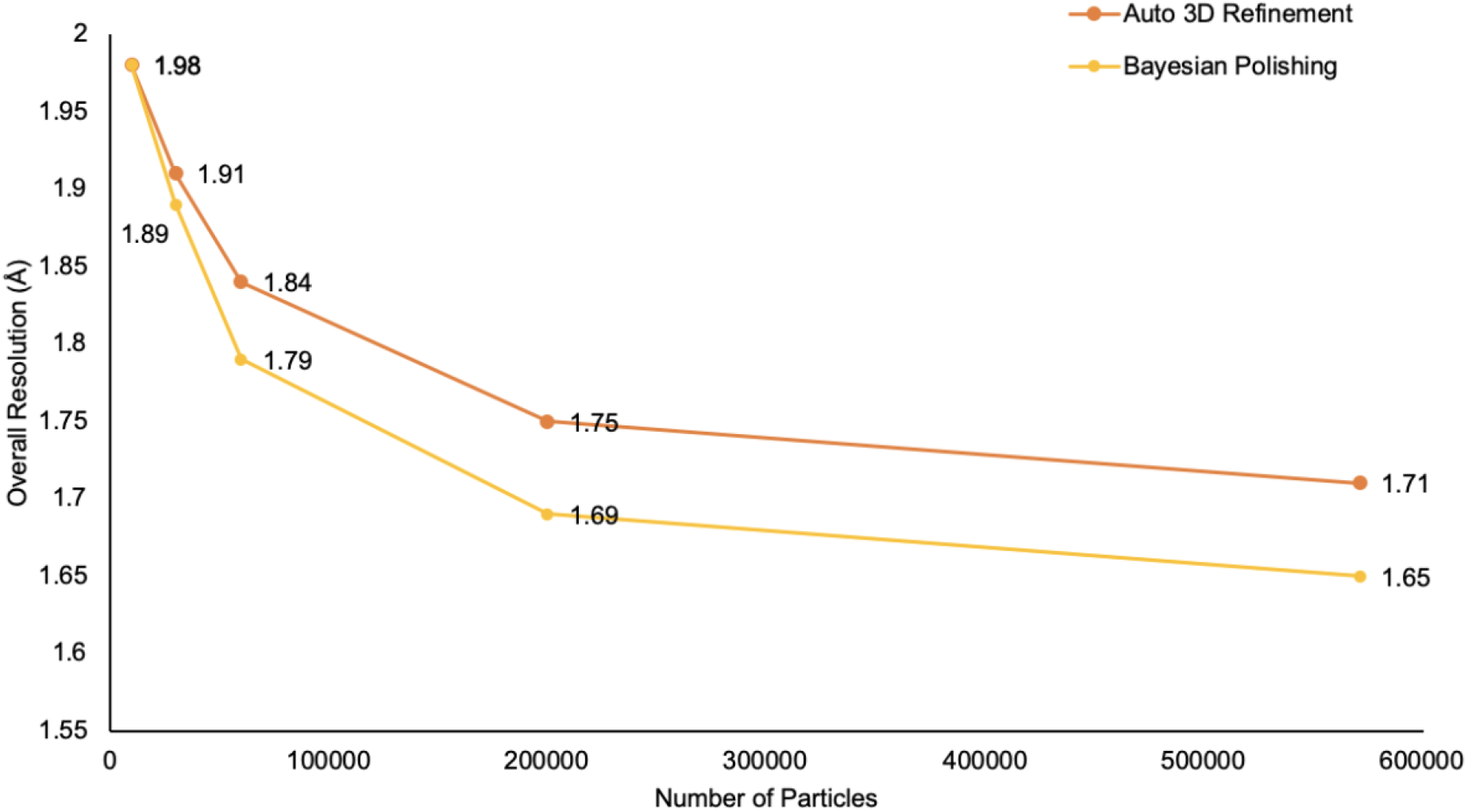
Resolution improvements from per-particle motion correction. To further improve the resolution of the apoferritin data sets collected using 9-hole image shift, Bayesian polishing was performed to further optimize per-particle beam-induced motion tracks, followed by another round of auto-refinement (Zivanov *et al*., 2019). All resolutions are reported in absolute angstrom and percentage of physical Nyquist frequency calculated from gold standard FSC estimates (Chen *et al*., 2013; Rosenthal & Henderson, 2003). Results are plotted as a function of percent of physical Nyquist frequency versus particle number and achievable resolutions are labelled.

**Figure S5.**
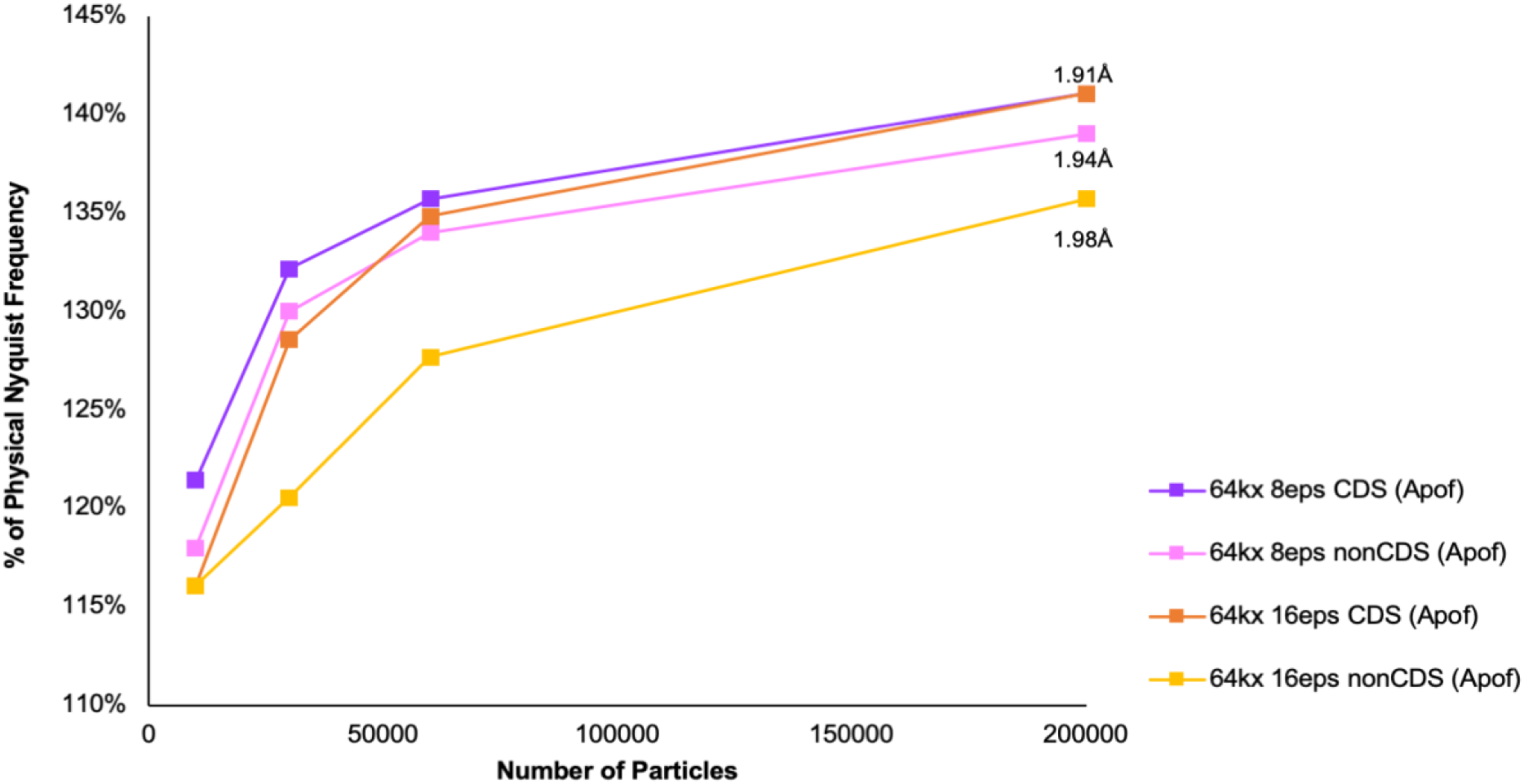
Balancing the effects of dose rate, coincidence loss, and noise reduction when imaging in CDS and nonCDS mode for apoferritin. Comparison of apoferritin data sets collected at 64,000x magnification with different dose rates. Results are plotted as a function of percent of physical Nyquist frequency versus particle number and highest achievable resolutions are labelled. All resolutions are reported in absolute angstrom and percentage of physical Nyquist frequency calculated from gold standard FSC estimates (Chen *et al*., 2013; Rosenthal & Henderson, 2003).

**Figure S6.**
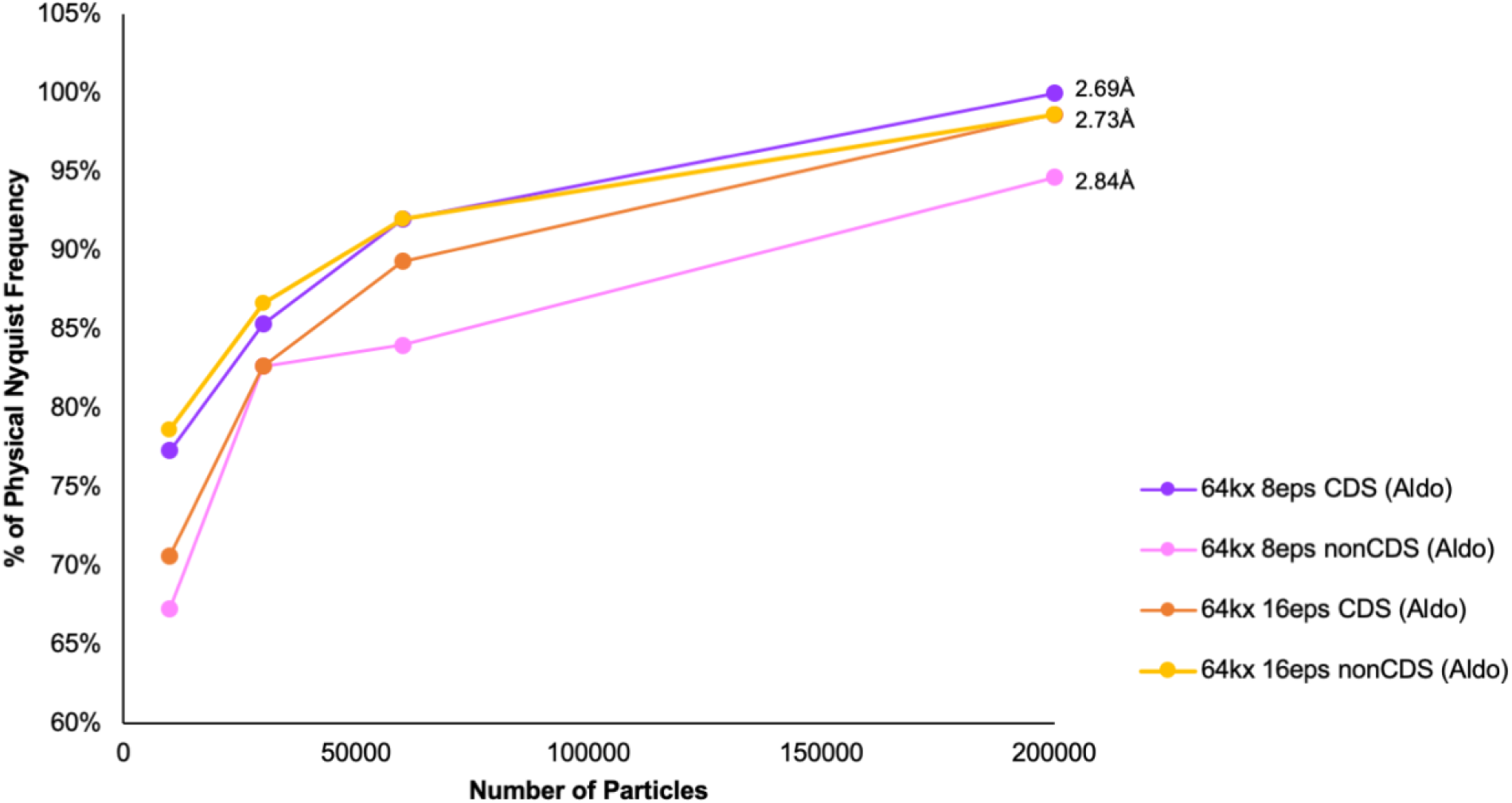
Balancing the effects of dose rate, coincidence loss, and noise reduction when imaging in CDS and nonCDS mode for aldolase. Comparison of aldolase data sets collected at 64,000x magnification. These results are plotted as a function of percent of physical Nyquist frequency versus particle number and highest achievable resolutions are labelled. All resolutions are reported in absolute angstrom and percentage of physical Nyquist frequency calculated from gold standard FSC estimates (Chen *et al*., 2013; Rosenthal & Henderson, 2003).

**Figure S7.**
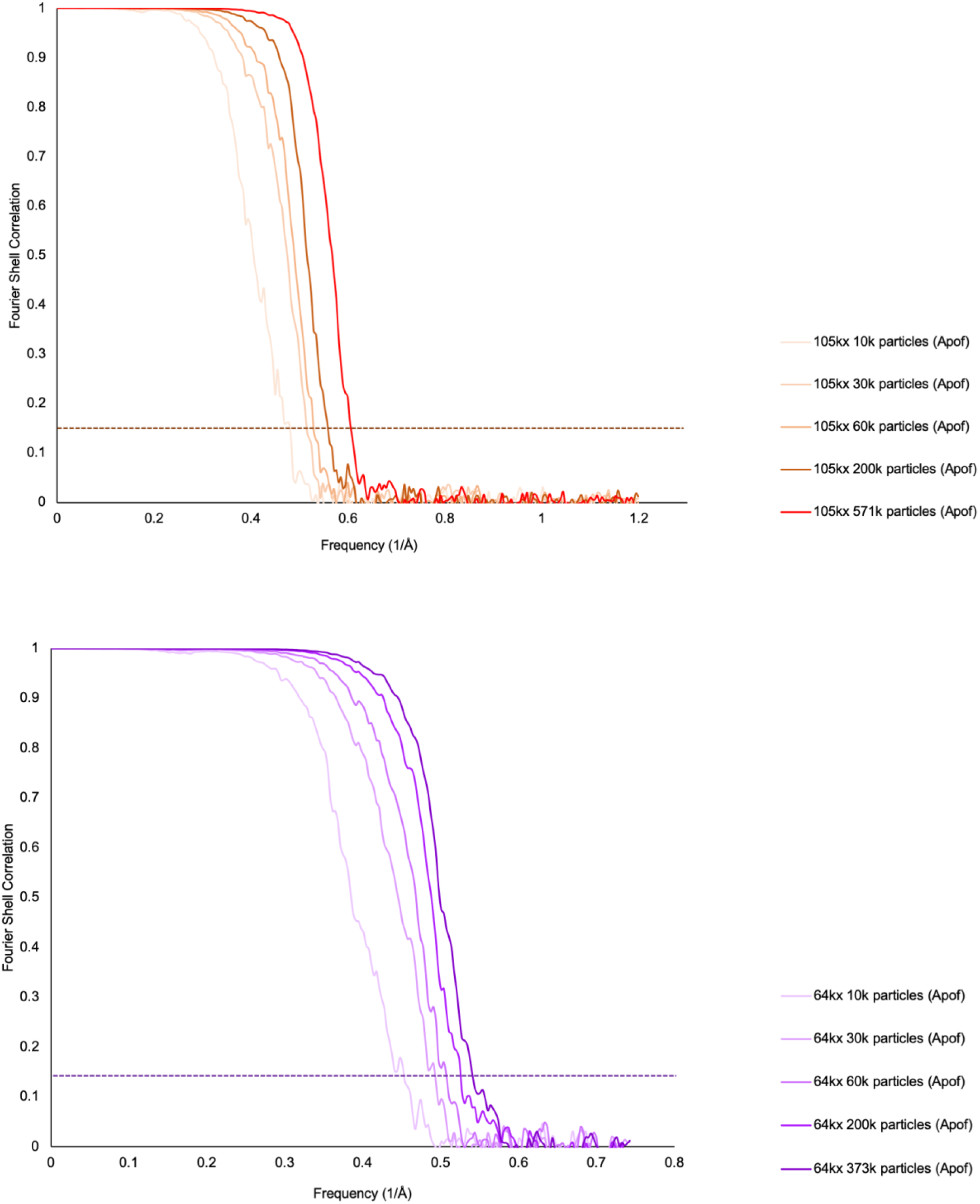
**Gold-standard FSC curves for the apoferritin reconstructions** from data subsets collected at 105kx (upper panel) and 64kx (lower panel).

## Notes

### Competing Interest Statement

The authors have declared no competing interest.

